# Development and qualification of an enzyme-linked immunosorbent assay to detect human serum immunoglobulin G reactive to multiple lineages of Lassa virus nucleoprotein

**DOI:** 10.64898/2025.12.25.696537

**Authors:** Heejin Yun, Faith Sigei, Nana Yaa Asiedua Appiah, Charlotte Naa Odey Quaye, Cecilia Adwoa Biaa Yankey, Eric Kyei-Baafour, Peter Hayes, Arianna Marini, Robert T Bailer, Marija Zaric, Kwadwo Asamoah Kusi

**Affiliations:** IAVI, New York, United States of America; Noguchi Memorial Institute for Medical Research, University of Ghana, Accra, Ghana

## Abstract

Lassa fever is a severe, often fatal febrile illness endemic to West Africa caused by Lassa virus (LASV). The viral nucleoprotein (NP) is a target antigen for serological assays to identify previous exposure to LASV. To our knowledge, there is no commercially available assay that reliably quantifies anti-LASV-NP IgG antibodies in human serum.

We report the development and qualification of an ELISA designed to detect and quantify anti-LASV-NP IgG in human serum samples. The assay employs recombinant Lineage IV LASV-NP immobilized on microwells to capture NP-specific IgG antibodies, which are then detected using horseradish peroxidase-conjugated anti-human IgG followed by TMB substrate and reading of optical densities. Optimal assay reagent concentrations, incubation times and temperature were determined along with assay positivity criteria and dynamic range. A reference standard prepared from pooled sera from donors in endemic Lassa fever regions was established and calibrated to the first WHO international standard for LASV antibodies. High and low positive controls for assay quality control were generated. Naïve human serum was used as a negative control. Following assay optimization, performance was assessed through assay qualification. The assay positivity criteria, lower limit of detection, upper and lower limits of quantitation, inter-assay precision, selectivity and dilutional linearity were determined.

Our anti-LASV-NP IgG ELISA was shown to reliably measure anti-LASV-NP IgG levels in human serum. The method demonstrated sensitivity, intra- and inter-assay precision, dilution linearity across its analytical range and specificity for anti-LASV-NP IgG. Establishing this assay represents an essential step toward strengthening LASV epidemiology research and supporting urgently needed development of a vaccine to prevent Lassa Fever.

## Introduction

Lassa fever (LF) is a severe, often fatal febrile illness caused by one of at least seven phylogenetic lineages of Lassa virus (LASV), a member of the *Arenaviridae* family [1–3]. LASV is endemic to West Africa [4] but also has pandemic potential. LASV is a zoonotic infection, the predominant host being the multimammate rat *Mastomys natalensis*, with transmission to humans via the gastrointestinal, respiratory or skin abrasion routes following contact with rodent urine or feces. Human to human transmission, including nosocomial outbreaks, may also occur via contact with blood, bodily fluids or excreta with virus excreted in urine for up to nine weeks [5]. There is currently no approved LF vaccine and LF is a WHO priority disease due to public health risk [6]. IAVI has been developing a vaccine candidate targeting LASV surface glycoprotein complex (GPC) encoded in a replication competent recombinant Vesicular Stomatitis Virus platform (rVSVΔG-LASV-GPC)[7]. The rVSVΔG platform has been successfully applied by Merck & Co., Inc., USA in development of a Zaire Ebola virus vaccine (ERVEBO) that has been approved by the FDA, EMA and African countries [8–12]. rVSVΔG-LASV-GPC vaccine candidate has been tested in a Phase 1 clinical trial (IAVI C102 study: NCT04794218) [13] and is under evaluation in a Phase 2a study (IAVI C105 study: NCT05868733), across West African clinical trial sites in Liberia, Ghana and Nigeria. Such studies aim to evaluate vaccine candidate safety and immunogenicity, thereby informing on vaccine design and further evaluation in clinical trials. Vaccine candidate immunogenicity is evaluated by assessments of humoral and cellular immune responses to LASV-GPC prior to and following vaccine candidate administration.

Since LF is endemic in countries that host current and future trial centers, it is important to assess participants in clinical trials of LASV vaccine candidates for prior exposure to LASV, enabling evaluation and understanding of the vaccine candidates’ safety and immunogenicity in the context of pre-existing immune responses to LASV. Nucleoprotein (NP) is one of the five major proteins encoded by LASV [14], it is not a component of current or planned vaccine candidate regimens and can be used as a target antigen in serological assessments to identify exposure to LASV. A commercial kit detecting anti-LASV-NP antibodies in human serum (ReLASV® Pan-Lassa NP IgG/IgM ELISA Kit, Zalgen Laboratory) was employed in IAVI C102 clinical trial to identify trial participants with pre-existing antibody responses to LASV-NP and thereby previous exposure to LASV [13]. However, these kits are no longer available, requiring development and qualification of a new enzyme-linked immunosorbent assay (ELISA) for detection of anti-LASV-NP antibodies. Development work was conducted at the Noguchi Memorial Institute for Medical Research (NMIMR), Accra, Ghana. The principle of the assay was as follows: anti-LASV-NP antibodies in samples reacted with recombinant LASV-NP antigen immobilized on ELISA plate microwells. Any immunoglobulin G (IgG) antibodies bound to LASV-NP were detected with horseradish peroxidase (HRP)-conjugated anti-human IgG antibodies, utilizing 3,3’,5,5’-tetramethylbenzidine (TMB) as HRP substrate with optical density (OD) values measured at 450nm with 620-650nm as reference. An anti-LASV-NP IgG reference standard curve and positive and negative controls were included on each assay plate, allowing detection and quantification of anti-LASV-NP IgG.

Several lineages of LASV are endemic to West Africa with lineages I, II and III predominating in Nigeria and lineage IV in Liberia and Sierra Leone [1] with LASV vaccine development approaches to date being focused on lineage IV [5]. There is a high level of protein sequence similarity between lineages with lineage IV LASV-NP sharing 90.5%, 89.6% and 91.6% amino acid identities with lineages I, II and III LASV-NP respectively [5], with lineage IV strains spreading across West Africa originating from ancestral Nigerian strains [15,16]. Therefore, to reduce cost and required volunteer sample volumes and increase assay throughput, ELISA development assessed the feasibility of detecting antibodies to multiple LASV-NP lineages with a single ELISA format.

The overall aim of the work was to develop and qualify an ELISA to detect anti-LASV-NP IgG antibodies in human serum, allowing assessment of participants in clinical trials of LASV vaccine candidates for potential pre and in-trial LASV exposure. The present report describes the successful development and qualification of such an assay, which was found to be reproducible and allowed for quantification of levels of IgG specific for multiple lineages of LASV-NP with one ELISA.

## Methods

### Study approval and informed consent

C105 clinical trial study protocol (NCT05868733) was approved by the institutional review boards at clinical trial sites in Nigeria, Liberia and Ghana. Participants provided written informed consent and completed an informed consent comprehension assessment prior to enrollment. The study started on 6^th^ March 2024 and is ongoing at the present time (December 2025). The study groups include adults, adolescents (twelve to seventeen years of age) and children (eighteen months to five years of age and six to eleven years of age) with informed consent obtained from the parents or guardians of non-adult participants.

### Blood sample collection and serum processing

Clinical trial blood samples were processed within six hours of collection. Serum tubes were centrifuged at 1,200 x*g* for 10 minutes and serum aliquots stored at -80°C.

### Enzyme-linked immunosorbent assay to detect anti-LASV-NP IgG

High-binding, flat-bottom EIA/RIA plates (Corning Incorporated, item 3590) were coated with soluble LASV-NP antigens: lineage II (Zalgen Labs, item LASV-R-0021), lineage III (Zalgen Labs, item LASV-R-0031) or lineage IV (Zalgen Labs, item LASV-R-0041) and stored at 4°C for 16-20 hours. Initial assay development assessed optimal ELISA signal with ELISA plate wells coated with different combinations of lineage II, III and IV LASV-NP at either 1 or 2 μg/mL total protein concentration in phosphate-buffered saline (PBS, Sigma-Aldrich, item D8537). After coating, plates were washed four times with 300 µL PBS with 0.05% v/v tween (PBS-T, Sigma-Aldrich, item P3563), with this washing procedure employed between subsequent incubations with test materials and development reagents. Plates were blocked with 200 µL PBS with 1% w/v casein (ThermoFisher Scientific, item 37582) for 1 hour. All incubations other than plate-coating were conducted initially at 37°C.

During assay development, a pool of anti-LASV-NP IgG positive serum samples identified from C105 trial participants was calibrated against a First WHO International Standard of anti-Lassa fever virus antibodies (NIBSC code 20/202) and was used to generate a standard curve and assay high (HPC) and low positive controls (LPC). Positive controls, negative control of a commercially supplied serum (Sigma-Aldrich, item H3667) from healthy humans not exposed to LASV, reagent blank wells (PBS 1% w/v casein) and test samples were included in triplicate in each assay plate and incubated for one hour. HRP-labelled anti-human IgG detection antibody (Sigma, item A0170) diluted 1:10,000 was added for one hour. Plates were developed by adding KPL Sureblue TMB 1-component peroxidase substrate (3,3’,5,5’–tetramethylbenzidine (TMB) SeraCare, item 5120-0074) for 20 minutes and the reaction stopped using TMB Stop Solution (Microimmune, item MI20031). Optical densities were read within five minutes using a Biotek 800TS microplate plate reader (450 nm with reference 620-650 nm). Optimal detection antibody concentration along with incubation times and temperatures for samples, detection antibody and HRP TMB substrate were assessed prior to assay qualification.

### Statistical analysis

Data were analyzed using GraphPad Prism version 10.6.1 (892) for Windows, GraphPad Software, San Diego, USA. A two-tailed non-parametric Wilcoxon matched pairs signed rank test assessed significant differences between two sets of paired values. A non-parametric Friedman test assessed significant differences within multiple groups of paired data. A two-tailed non-parametric Spearman test computed correlation coefficients between two variables. The threshold for significance was defined as p < 0.05.

## Results and discussion

### Anti-LASV-NP IgG ELISA development and qualification

#### Initial assay establishment

Initial assay development assessed optimal ELISA signal (OD_450nm_) with ELISA plate wells coated with either a combination of lineage II, III and IV (in 1:1:1 ratio) or lineage IV alone LASV-NP at 1 or 2 μg/mL total protein concentration. A WHO International Standard of anti-Lassa fever virus antibodies (First WHO International Standard for anti-Lassa fever virus antibodies, NIBSC code 20/202) was used to generate a standard curve, starting at concentration of anti-LASV-NP IgG of 50 IU/mL and followed by nine subsequent two-fold serial dilutions in 1X PBS w/v 1% casein. Positive control (First WHO International Standard for anti-Lassa fever virus antibodies diluted to WHO-assigned 10 IU/mL), negative control and reagent blank wells were included in triplicate in each assay plate. Incubations and signal development were conducted as described above. Fig 1 displays mean ELISA signal values plotted against WHO-assigned international units (IU) for anti-NP antibody content of the First WHO International Standard for anti-Lassa fever virus antibodies. The concentration of anti-LASV-NP IgG in samples was extrapolated by four-paramenter logistic (4PL) standard curves using Biotek GEN5 version 3.16 Software, Microsoft Excel and GraphPad Prism. S1 Table displays information regarding standard curve OD and interpolated positive control values. The ELISA utilizing lineage IV LASV-NP coated at 2 µg/mL was selected for final assay qualification as this resulted in the best curve fit, an R-squared value > 0.98 at 0.996, low negative control background signal, a wide signal span across the standard curve and a positive control recovery value similar to the nominal expected value at 89% recovery (S2 Table).

**Fig 1.**
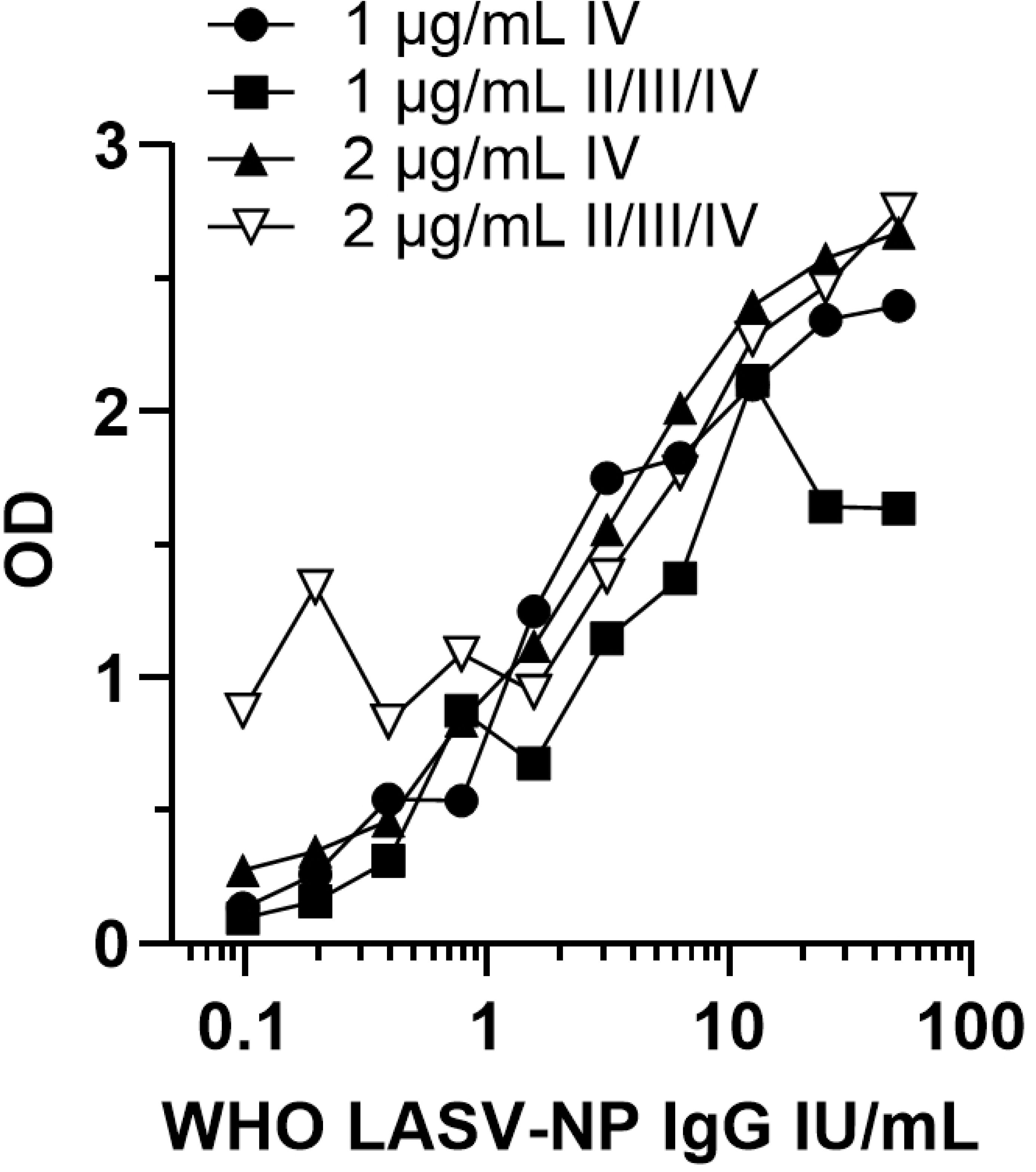
LASV-NP antigen lineage and coating concentration. A combination of LASV-NP lineage II/III/IV or lineage IV alone as coating antigens at either 1 or 2 μg/mL were assessed against a standard curve using the First WHO International Standard for anti-Lassa fever virus antibodies, starting at 50 IU/mL with serial two-fold dilutions. OD values are plotted against WHO-assigned international units (IU) for anti-NP antibody content of the standard. Closed circles: 1 μg/mL lineage IV, closed squares: 1 μg/mL lineages II/III/IV, closed triangles: 2 μg/mL lineage IV and open inverted triangles: 2 μg/mL lineages II/III/IV.

#### Generation of a new anti-LASV-NP IgG reference serum pool

The First WHO International Standard for anti-Lassa fever virus antibodies is available in limited quantities through NIBSC. Therefore, a new reference standard material was produced using anti-LASV-NP IgG positive serum samples from C105 clinical trial participants for further assay development and ultimate application to clinical trial testing, with calibration against the WHO reference standard. Pre-vaccination serum samples from forty-four C105 clinical trial participants that tested positive in an ELISA [13] to detect anti-LASV-GPC IgG antibodies, were tested at 1:100 dilution in the anti-LASV-NP IgG ELISA using the conditions selected above (LASV lineage IV). Fig 2 (left panel) displays anti-LASV-NP IgG ELISA OD values for these samples and assay controls. To provide additional reference material, post-vaccination serum samples from the five participants who had the highest pre-vaccination OD values were assessed using the anti-LASV-NP IgG ELISA. All post-vaccination samples tested resulted in high OD values (Fig 2 right panel). Serum samples donated by the four trial participants with the highest OD values were pooled to provide a new reference for subsequent assay development. The concentration in IU/mL of this new reference serum pool was assessed in assays utilizing both LASV-NP lineages II, III and IV and IV alone as coating antigens by comparison of serially diluted new reference serum pool and WHO reference standard. At this stage of assay development, the new reference serum pool was assigned an interim concentration of 418.0 and 544.8 IU/mL for LASV-NP lineages II, III and IV and lineage IV alone, respectively (S2 Table).

**Fig 2.**
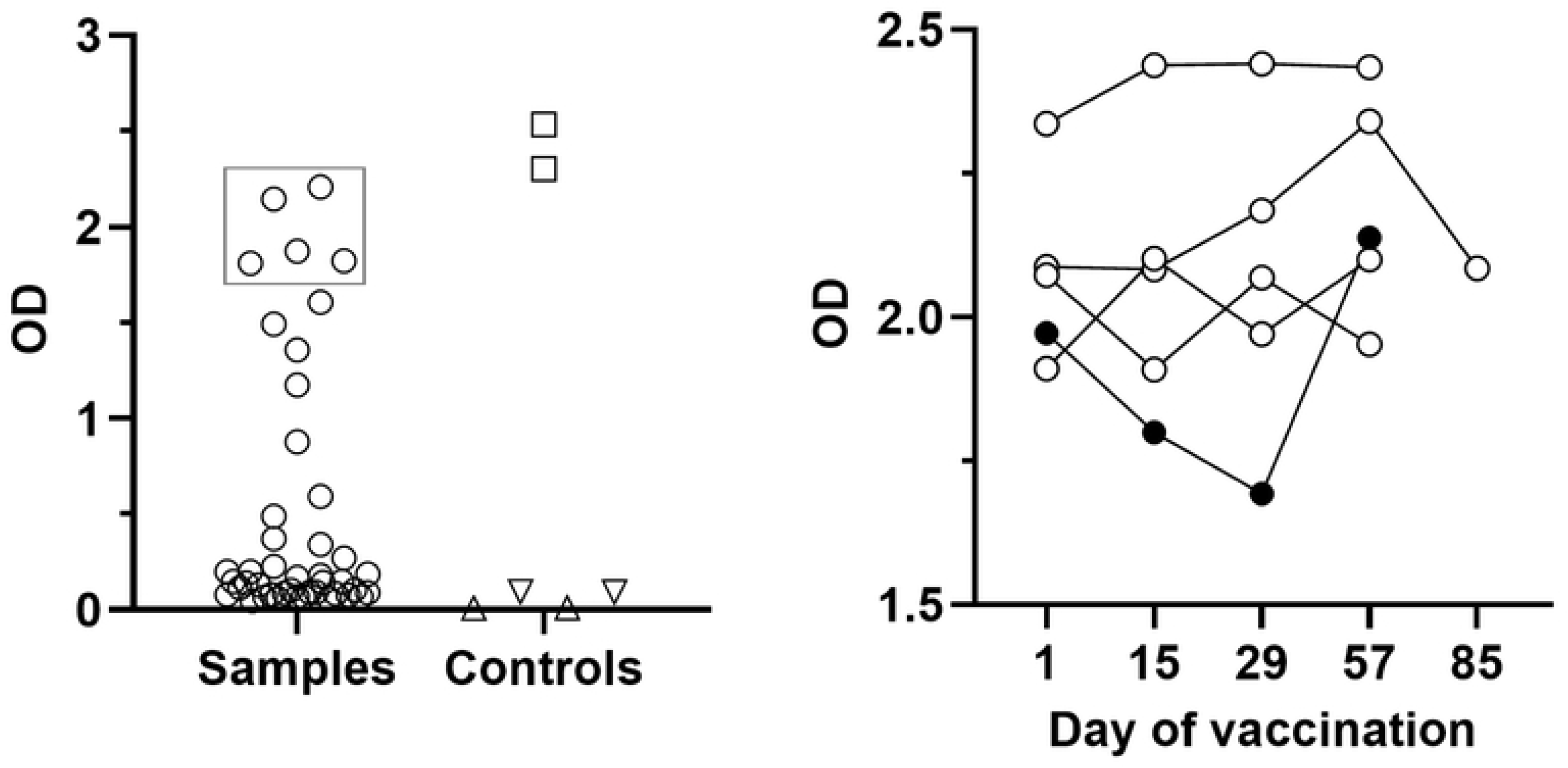
Generation of a new reference serum pool. **Left panel:** Anti-LASV-NP IgG ELISA OD values for C105 clinical trial serum samples testing positive for anti-LASV-GPC IgG prior to vaccination (circles). Additional samples from five trial participants with the highest OD values (within the box) were selected for further testing. OD values for ELISA positive controls (squares), negative controls (inverted triangles) and blank wells (triangles) are displayed. **Right panel:** Anti-LASV-NP IgG ELISA OD values for serum samples from the five selected trial participants re-tested at pre-vaccination (day 1) and post-vaccination (days 15 to 85) time points. Samples from four trial participants (open circles) were pooled to provide a new reference serum standard.

#### Cross lineage detection of anti-LASV-NP IgG

Given that LASV lineages I, II and III predominate in Nigeria, whilst lineage IV circulates in Liberia and Sierra Leone [1], it would be desirable to have one assay to detect anti-LASV-NP IgG in serum samples from all these countries. Anti-LASV-NP IgG ELISA OD signals were assessed in ten pre-vaccination anti-LASV-GPC IgG ELISA-positive C105 clinical trial samples from Nigeria and Liberia and a WHO international reference panel of seven samples for anti-LASV antibodies (21/332) from Nigeria and Sierra Leone. Two coating antigen lineage strategies were compared: LASV lineage IV NP only and combined LASV lineages II/III/IV NPs (1:1:1 ratio), both conditions were coated at 2 µg/mL total protein antigen concentration. All plates included controls and standard curves generated using the new reference serum pool. Fig 3 displays ELISA signal values obtained with these samples and coating antigens.

**Fig 3.**
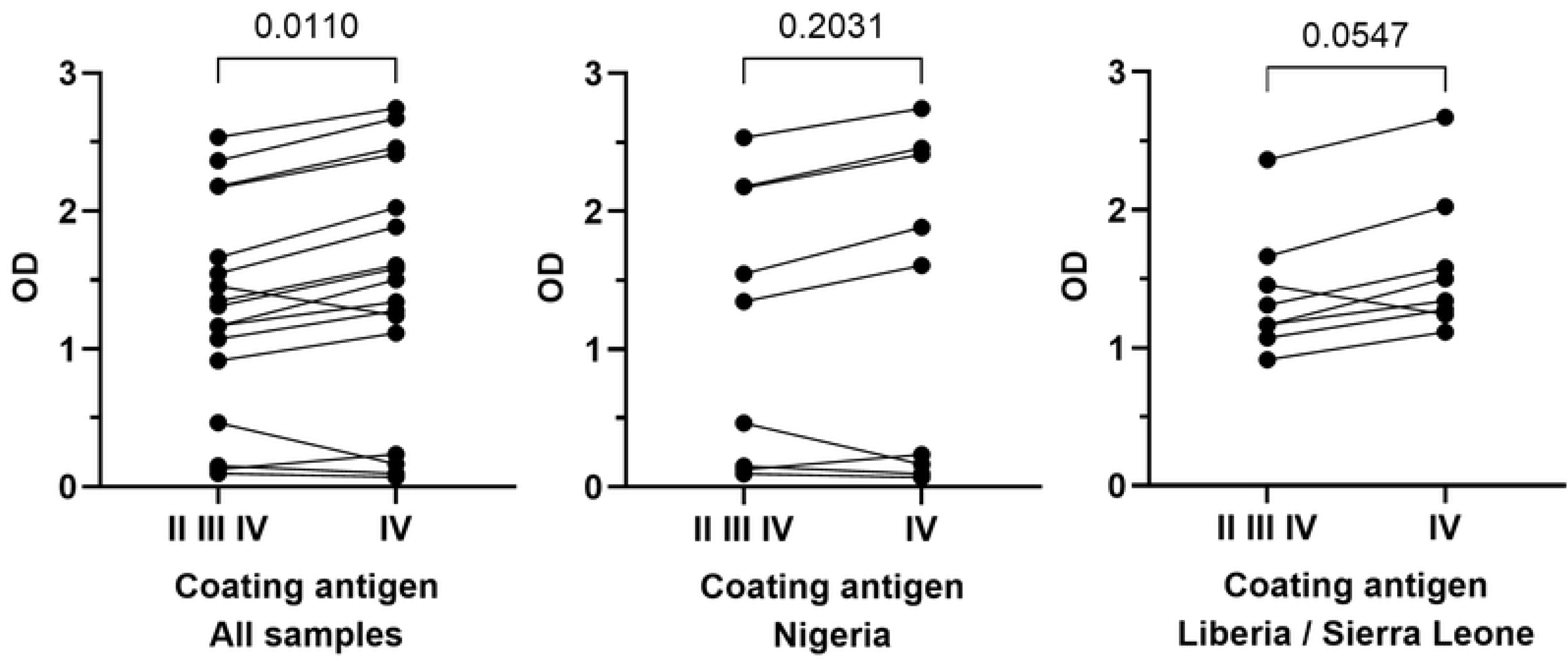
Evaluation of LASV-NP coating antigen lineages. **Left panel (all samples):** Anti-LASV-NP IgG ELISA OD values for all samples tested (WHO panel and C105 clinical trial serum samples testing positive for anti-LASV GPC IgG prior to vaccination) using either combined LASV-NP lineage II, III and IV or lineage IV alone as ELISA coating antigens. These data are segregated into samples from Nigeria (**middle panel**) where LASV lineages I, II and III predominate or Liberia and Sierra Leone (**right panel**) where LASV lineage IV predominates. The p values of Wilcoxon matched pairs signed rank tests between two datasets are displayed above the datapoints.

Both coating antigen lineage conditions generated ELISA signals from samples derived from all geographic locations, demonstrating that either coating antigen lineage condition can detect anti-LASV-NP IgG antibodies in samples where either LASV lineages I, II and III or lineage IV predominate. The OD signal values were marginally but significantly higher with lineage IV coating antigen compared with combined lineage II, III, and IV (median OD of 1.50 and 1.31 respectively, p = 0.0110, Wilcoxon matched pairs signed rank test, Fig 3 left panel). Samples derived from Nigeria (predominant LASV lineages I, II and III) generated signals with both coating antigen conditions with a trend to higher signals using lineage IV NP protein (median OD lineage IV and II, III and IV of 1.61 and 1.35 respectively, p = 0.2031, Fig 3 middle panel). Similarly, Liberia or Sierra Leone samples (predominant LASV lineage IV) also generated signals with both coating antigen conditions with a trend to higher signals using lineage IV NP protein (median OD lineage IV and II, III and IV of 1.42 and 1.24 respectively, p = 0.0547, Fig 3 right panel). Coating antigen lineage IV (2 μg/mL) was chosen for further assay development as this resulted in optimal ELISA signal for samples from Nigeria, Sierra Leone and Liberia.

#### Evaluation of ELISA incubation parameters

The following ELISA incubation parameters were evaluated using lineage IV LASV-NP at 2 μg/mL as coating antigen: dilution (1:5,000 or 1:10,000) of anti-human IgG detection antibody, sample (60 or 120 minutes), detection antibody (60 or 120 minutes) and TMB substrate (10 or 20 minutes) incubation times and assay incubation temperature (room temperature or 37°C). OD values of serially diluted new reference pool are displayed in Fig 4 (left panel) for the different assay parameters. Initial assessment of assay parameters indicated better signals in terms of range of OD values across the standard curves and lower negative control OD values with 20 minutes TMB substrate incubation time. ELISA parameters of detection antibody dilution, sample incubation time and assay incubation temperature were further investigated in 10 different negative control samples with the aim of minimizing background signal (Fig 4 right panel). Based on higher specific and lower background signals the optimal ELISA parameters were chosen of 60-minute sample and detection antibody incubation times with detection antibody diluted 1:10,000, 20-minute incubation of TMB substrate with the ELISA conducted at 37°C.

**Fig 4.**
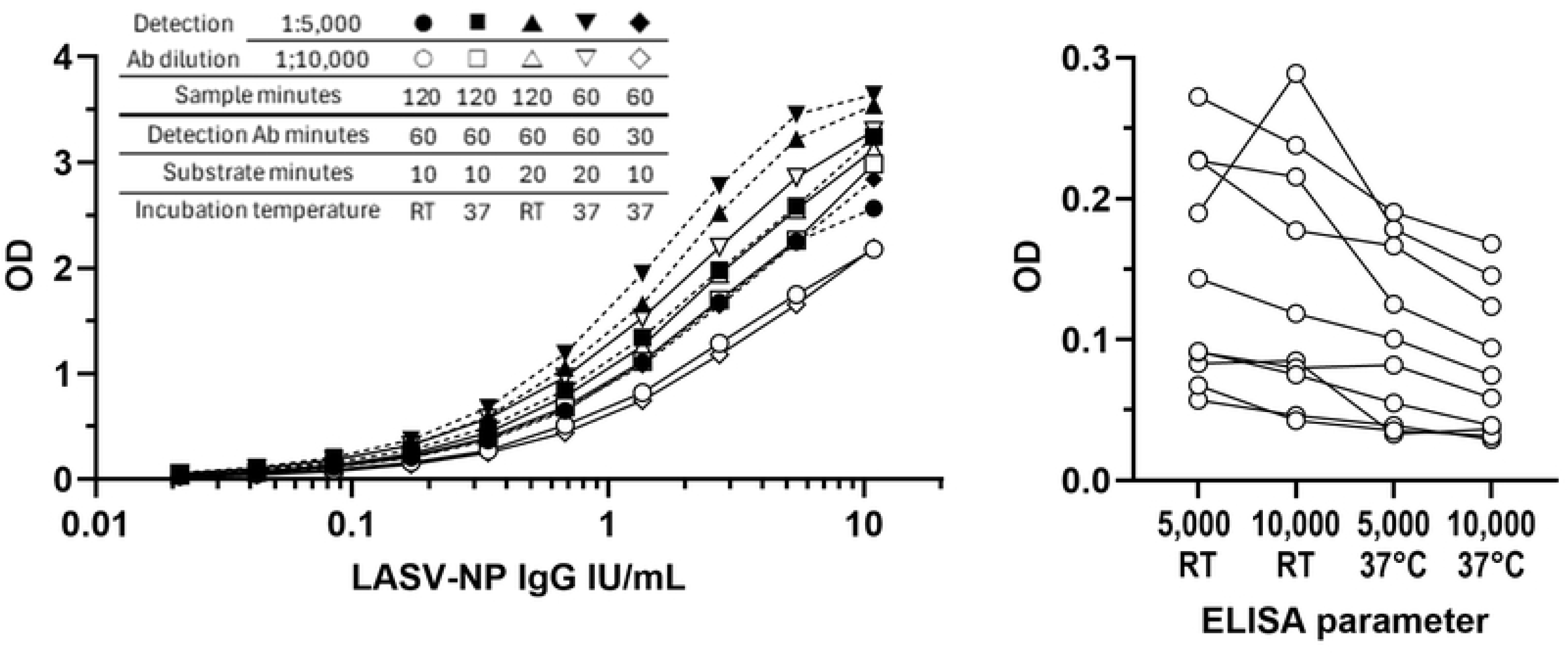
Optimization of assay conditions. **Left panel:** ELISA OD values under different assay conditions. Detection antibody dilutions are indicated as 1:5,000 (closed data points) and 1:10,000 (open data points). Assay parameters of time of sample, detection antibody and TMB substrate incubation and incubation temperature are respectively indicated by circles: 120 minutes, 60 minutes, 10 minutes, room temperature, squares: 120 minutes, 60 minutes, 10 minutes, 37°C, triangles: 120 minutes, 60 minutes, 20 minutes, room temperature, inverted triangles: 60 minutes, 60 minutes, 20 minutes, 37°C and diamonds: 60 minutes, 30 minutes, 10 minutes, 37°C. **Right panel:** ELISA OD values for 10 LASV negative serum samples tested with either detection antibody diluted at 1:5,000 or 1:10,000 and incubated at room temperature (RT) or 37°C. Ab: antibody.

Using the new ELISA parameters with LASV-NP lineage IV as coating antigen, the new reference serum pool was assigned a concentration of 464.6 IU/mL by comparison of serially diluted new reference serum pool and WHO reference standard.

#### Assay positivity criteria

Serum samples from sixty-three healthy Ghanaian volunteers with no history of LF were tested using the LASV-NP IV lineage ELISA using the newly established assay parameters. Each sample was tested by three operators: each operator testing every sample three times. Two outlier samples with mean ODs of 2.99 and 0.226 were identified and excluded using the ROUT test (Q = 1%). OD values for the remaining sixty-one samples are displayed in Fig 5 and S3 Table. The assay positivity point was determined to be an OD of 0.182 by calculating the mean OD (0.071) of these values and adding three standard deviations (3 x 0.037). Therefore, samples yielding OD values of ≤0.182 were defined as anti-LASV-NP IgG negative in this ELISA.

**Fig 5.**
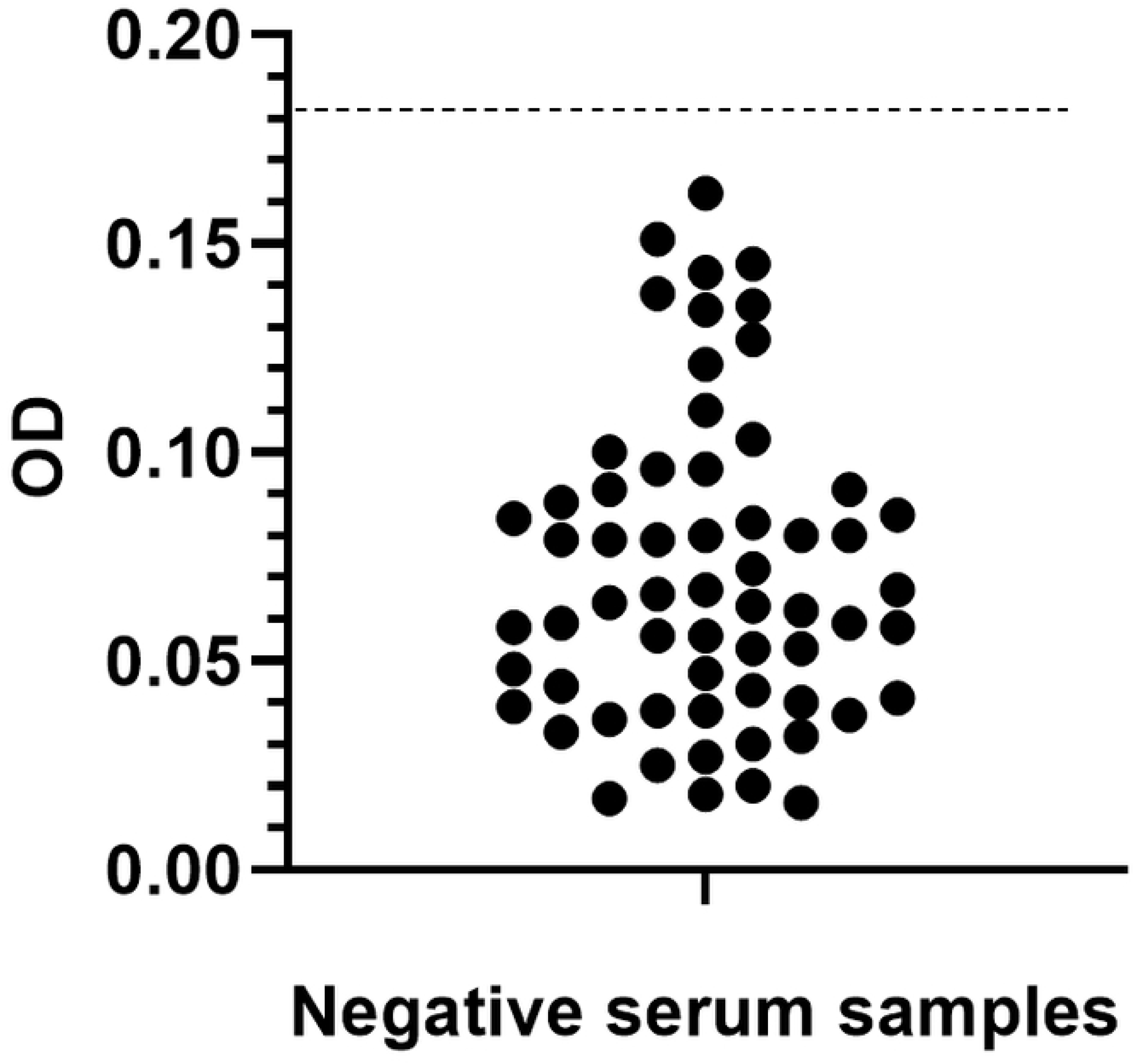
Determination of assay positivity point. ELISA OD values for serum samples from sixty-one healthy Ghanaian volunteers with no history of LF. The dotted line represents the assay positivity point OD value of 0.182.

#### Anti-LASV-NP IgG ELISA dynamic range and positive controls

The assay upper limit of quantification (ULOQ) and lower limit of quantification (LLOQ) were defined as the highest and lowest concentrations, respectively, at which the percent recovery between measured and expected values fell within the 75-125% recovery range and remained precise (CV <25%). ULOQ and LLOQ were assessed with a ten-point curve using a two-fold serial dilution series starting from a 1:60 dilution of the new reference serum pool in duplicate wells. The measured concentrations of each point of this curve were interpolated from a standard curve generated using two-fold serial dilutions of the reference serum pool starting at 1:50 dilution. Nine determinations were performed by three different operators, with three independent runs conducted on separate days allowing for assessment of assay precision (%CV). A low positive control (LPC) consisting of a 1:1000 dilution of the new reference serum pool was chosen to generate an anti-LASV-NP IgG value approximately four times above the LLOQ at 0.47 IU/mL. A lower dilution of 1:250 of the new reference serum pool was selected as a high positive control (HPC) below the ULOQ, equating to 1.86 IU/mL.

Data are displayed in Table 1. Dilution series ODs and interpolated IU/mL were precise (CV <25%) over the first seven dilutions equating to an ULOQ of 7.74 IU/mL and a LLOQ of 0.12 IU/mL, with the LLOQ resulting in OD values (mean 0.187) above the assay positivity point (OD 0.182). Recovery of interpolated IU/mL values over this range was between 94.7% and 99.6% compared with nominal values. Both HPC and LPC yielded precise IU/mL values with recoveries between 98.3% and 107.3% of expected values. Acceptable positive control recoveries of between 75% and 125% of expected values would equate between 0.349 to 0.581 and 1.394 to 2.323 anti-LASV-NP IgG IU/mL for LPC and HPC, respectively.

**Table 1.**
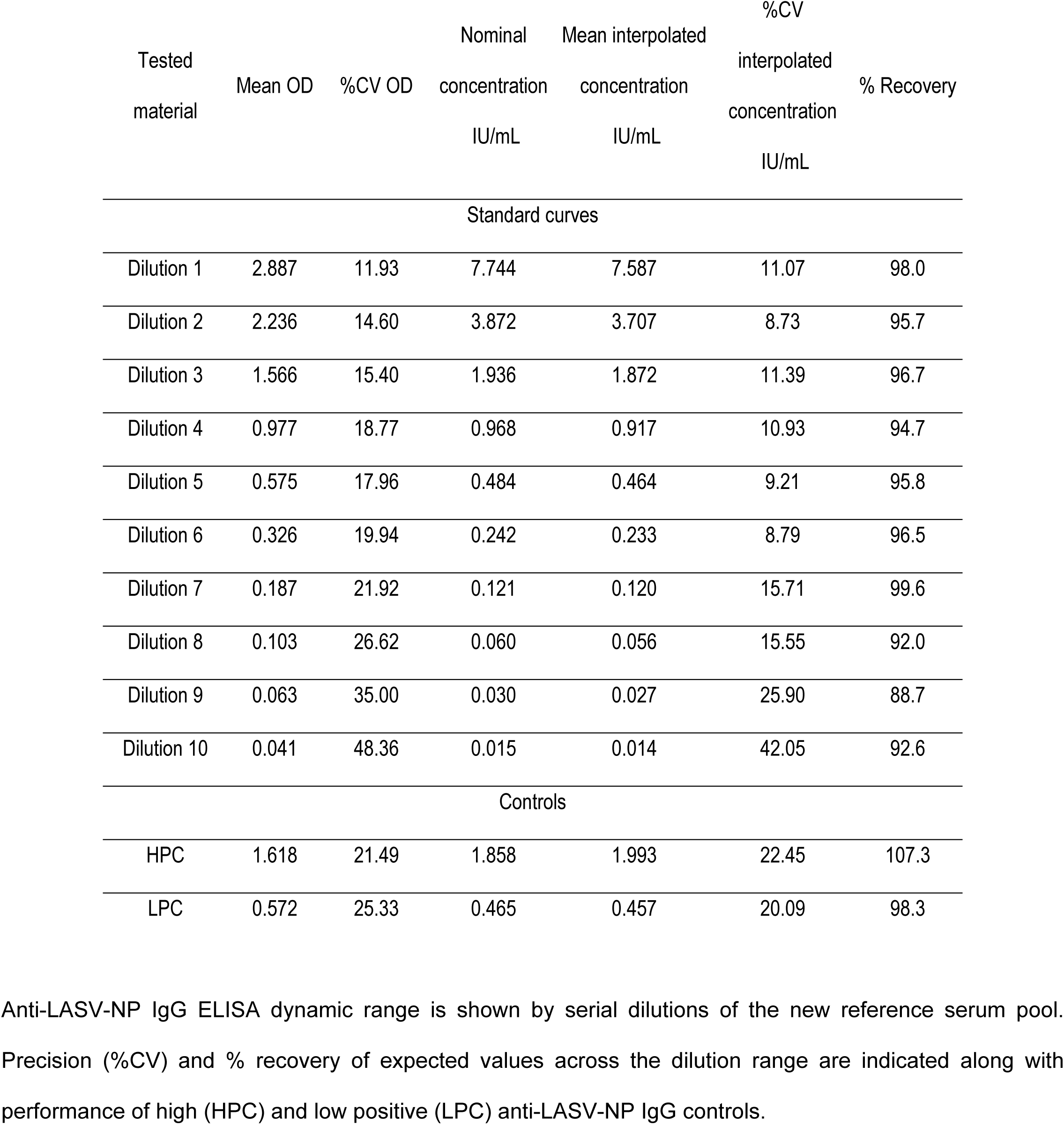
ELISA dynamic range and controls.

| Tested material | Mean OD | %CV OD | Nominal concentration<br>IU/mL | Mean interpolated concentration<br>IU/mL | %CV interpolated concentration<br>IU/mL | % Recovery |
| --- | --- | --- | --- | --- | --- | --- |
| Standard curves |  |  |  |  |  |  |
| Dilution 1 | 2.887 | 11.93 | 7.744 | 7.587 | 11.07 | 98.0 |
| Dilution 2 | 2.236 | 14.60 | 3.872 | 3.707 | 8.73 | 95.7 |
| Dilution 3 | 1.566 | 15.40 | 1.936 | 1.872 | 11.39 | 96.7 |
| Dilution 4 | 0.977 | 18.77 | 0.968 | 0.917 | 10.93 | 94.7 |
| Dilution 5 | 0.575 | 17.96 | 0.484 | 0.464 | 9.21 | 95.8 |
| Dilution 6 | 0.326 | 19.94 | 0.242 | 0.233 | 8.79 | 96.5 |
| Dilution 7 | 0.187 | 21.92 | 0.121 | 0.120 | 15.71 | 99.6 |
| Dilution 8 | 0.103 | 26.62 | 0.060 | 0.056 | 15.55 | 92.0 |
| Dilution 9 | 0.063 | 35.00 | 0.030 | 0.027 | 25.90 | 88.7 |
| Dilution 10 | 0.041 | 48.36 | 0.015 | 0.014 | 42.05 | 92.6 |
| Controls |  |  |  |  |  |  |
| HPC | 1.618 | 21.49 | 1.858 | 1.993 | 22.45 | 107.3 |
| LPC | 0.572 | 25.33 | 0.465 | 0.457 | 20.09 | 98.3 |

Anti-LASV-NP IgG ELISA dynamic range is shown by serial dilutions of the new reference serum pool. Precision (%CV) and % recovery of expected values across the dilution range are indicated along with performance of high (HPC) and low positive (LPC) anti-LASV-NP IgG controls.

#### Inter-assay precision and specificity

Assay precision describes the variability across repeated measures of the same test samples. Inter-assay precision was evaluated by testing twenty serum samples previously determined to be positive for anti-LASV-NP IgG responses consisting of WHO international panel (NIBSC) and C105 trial pre-vaccination participants’ samples. Each sample was tested nine times; three times each by three operators on separate occasions with %CV values determined across the replicates as a measure of assay precision. IU/mL values for the expected LASV-NP IgG positive samples were precise with seventeen of the twenty (85%) samples yielding CVs of less than 25% (Table 2). Two of the three samples with CVs >25% (C105-12 and C105-13 at 27.9% and 72.4%, respectively) were determined to have IU/mL values around the assay LLOQ (0.12 IU/mL) and therefore may be expected to yield more variable data.

**Table 2.** ELISA inter-assay precision.

| Sample ID | Operator 1 IU/mL |  |  | Operator 2 IU/mL |  |  | Operator 3 IU/mL |  |  | All operators |  |
| --- | --- | --- | --- | --- | --- | --- | --- | --- | --- | --- | --- |
|  | RUN 1 | RUN 2 | RUN 3 | RUN 1 | RUN 2 | RUN 3 | RUN 1 | RUN 2 | RUN 3 | Mean | %CV |
| NIBSC-20/226 | >ULOQ<sup>a</sup> | >ULOQ<sup>a</sup> | 11.38 | 12.13 | >ULOQ<sup>a</sup> | 12.61 | >ULOQ<sup>a</sup> | 13.64 | >ULOQ<sup>a</sup> | 12.46 | 9.11 |
| NIBSC-20/244 | 7.85 | >ULOQ<sup>a</sup> | 12.70 | 7.47 | 10.69 | 10.43 | 10.32 | 7.90 | 13.51 | 10.13 | 22.35 |
| NIBSC-20/228 | 8.84 | 8.51 | 9.44 | 7.62 | 10.96 | 9.15 | 9.16 | 8.49 | 7.75 | 8.88 | 11.24 |
| NIBSC-20/204 | 8.83 | 7.16 | 8.12 | 8.58 | 9.33 | 7.70 | 7.78 | 6.35 | 6.99 | 7.87 | 12.11 |
| C105-01 | 5.79 | 6.83 | 6.86 | 5.45 | 8.19 | 6.94 | 7.39 | 4.16 | 6.46 | 6.45 | 18.28 |
| C105-02 | 6.89 | 5.45 | 5.91 | 6.63 | 6.20 | 7.40 | 6.66 | 5.38 | 6.45 | 6.33 | 10.51 |
| C105-03 | 2.96 | 4.60 | 4.07 | 3.60 | 5.10 | 3.95 | 4.38 | 2.67 | 3.49 | 3.87 | 20.08 |
| NIBSC-20/222 | 3.72 | 3.56 | 3.91 | 3.97 | 3.59 | 3.51 | 3.82 | 3.74 | 2.83 | 3.63 | 9.27 |
| C105-04 | 2.36 | 2.61 | 3.59 | 3.67 | 3.74 | 3.81 | 3.34 | 2.27 | 2.33 | 3.08 | 21.75 |
| C105-05 | 2.40 | 3.53 | 2.78 | 2.54 | 3.08 | 2.97 | 3.70 | 3.30 | 2.70 | 3.00 | 14.83 |
| NIBSC-20/246 | 1.91 | 2.61 | 2.78 | 2.58 | 2.95 | 2.98 | 4.00 | 2.57 | 2.21 | 2.73 | 21.34 |
| C105-06 | 2.02 | 2.19 | 2.21 | 1.76 | 2.45 | 2.55 | 4.31 | 1.55 | 2.11 | 2.35 | 33.94 |
| C105-07 | 1.67 | 2.36 | 2.28 | 2.01 | 2.40 | 3.08 | 2.85 | 1.75 | 2.18 | 2.29 | 20.30 |
| C105-08 | 1.57 | 2.43 | 2.57 | 1.84 | 2.02 | 2.46 | 2.05 | 1.65 | 1.68 | 2.03 | 18.74 |
| C105-09 | 1.73 | 2.46 | 1.75 | 1.50 | 2.39 | 2.22 | 2.34 | 1.31 | 1.81 | 1.94 | 21.54 |
| C105-10 | 1.67 | 2.02 | 1.46 | 1.76 | 2.06 | 1.97 | 1.95 | 1.84 | 1.92 | 1.85 | 10.33 |
| NIBSC-20/248 | 1.32 | 1.47 | 1.87 | 1.59 | 2.03 | 2.18 | 1.90 | 1.74 | ND<sup>b</sup> | 1.77 | 16.45 |
| C105-11 | 0.64 | 0.60 | 0.77 | 0.91 | 0.96 | 0.83 | 1.10 | 0.82 | 0.71 | 0.81 | 19.43 |
| C105-12 | 0.12 | 0.51 | 0.13 | 0.09 | 0.18 | 0.13 | 0.12 | 0.13 | 0.17 | 0.18 | 72.39 |
| C105-13 | 0.09 | 0.08 | 0.08 | 0.05 | 0.12 | 0.07 | 0.07 | 0.06 | 0.10 | 0.08 | 27.87 |

Anti-LASV-NP IgG ELISA IU/mL values and precision (%CV) across nine assay tests for each sample evaluated by three operators. Samples are ranked from highest to lowest mean IU/mL values.
<sup>a</sup>ULOQ: upper limit of quantification. <sup>b</sup>ND: Not done.

Assay specificity was assessed for these twenty anti-LASV-NP IgG positive samples and the sixty-one negative samples from healthy Ghanaian volunteers with no history of LF previously used to determine assay positivity criteria (Fig 5). Each data value was assigned as positive (OD >0.182) or negative (OD ≤0.182) and the concordance determined between the nine tests across three operators for each sample. For the assay to be deemed to be specific, eight (88.9%) of the nine determinations for each sample must be concordant with this achieved in at least 80% of samples. This requirement was achieved for the expected negative samples with fifty-one of sixty-one (83.6%) samples being fully concordant in all nine determinations and 54 (88.5%) of these samples concordant in at least eight of nine determinations (S3 Table). All expected negative samples with concordance of less than eight of nine determinations had mean OD values considerably higher than the mean OD of the group (OD 0.071) between OD 0.121 and 0.162. Such values would be more likely to fall into either positive or negative determinations around the assay positivity point of OD 0.182. The requirement was also met for expected positive samples (S4 Table) with eighteen of twenty (90%) of samples being fully concordant. The two samples with less than eight of nine determinations being concordant yielded mean OD values of 0.12 and 0.25, close to the assay positivity point.

#### Dilution linearity

To demonstrate that assay values are not affected by sample dilution, assay dilution linearity was assessed by serially diluting three anti-LASV-NP IgG positive serum samples to span the range of the standard curve. Dilutions were tested independently by three operators and reported in IU/mL following adjustment for the dilution factor in Fig 6 and Table 3. Assay accuracy across the 1:100 to 1:1,600 dilutions tested was demonstrated by linear results for anti-LASV-NP IgG concentrations (IU/mL) across these dilutions, with CVs across the three operators being less than 25%. Simple linear regression of anti-LASV-NP IgG IU/mL values prior to dilution adjustment against the reciprocal of the serum dilution factor demonstrated dilution linearity with R-squared values of at least 0.9887 for eight of the nine determinations (Table 3).

**Fig 6.**
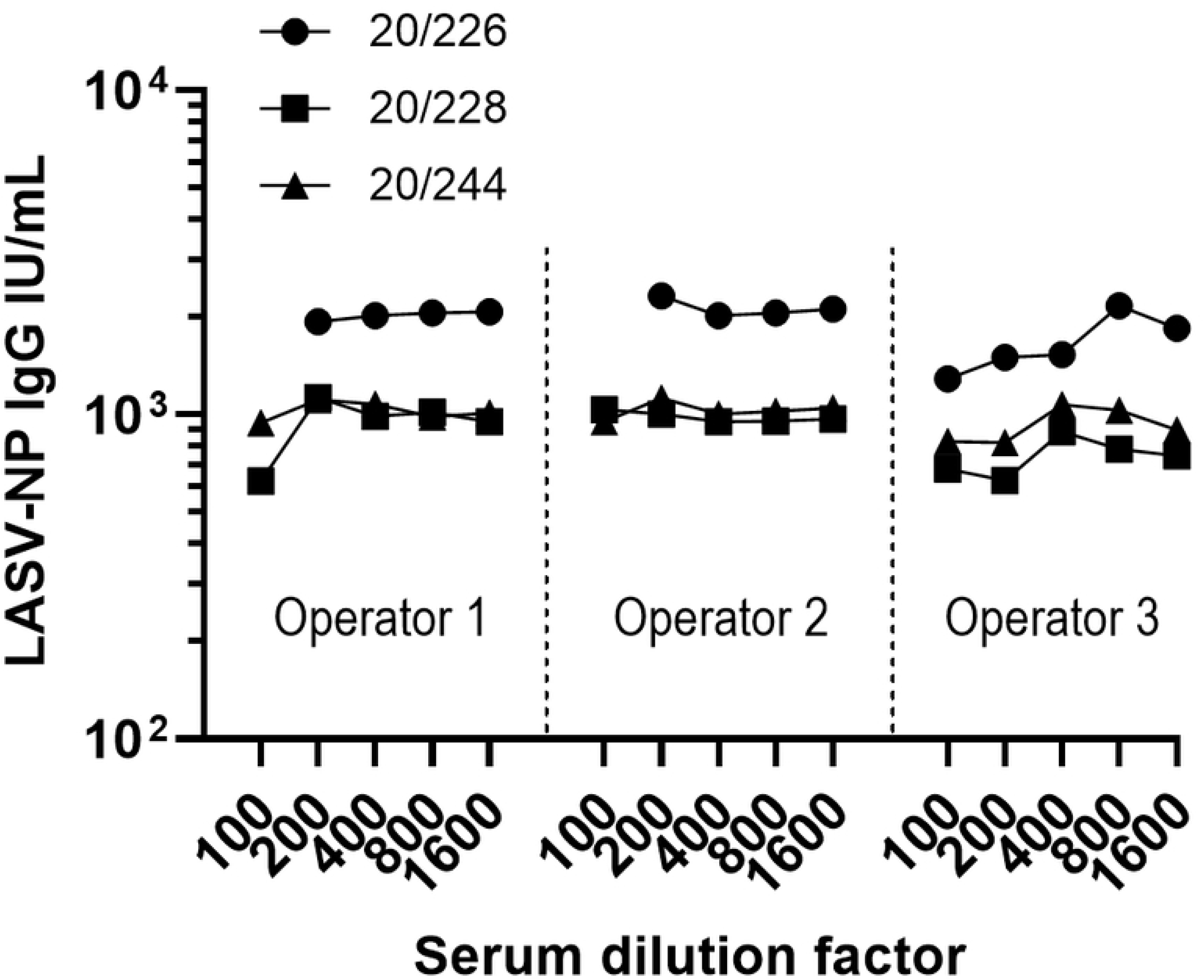
Assay dilution linearity. Dilution linearity of three anti-LASV-NP IgG positive samples across dilutions from 1:100 to 1:1,600 each tested by three operators. Sample 1 (NIBSC 20/226, circle), sample 2 (NIBSC 20/228, square) and sample 3 (NIBSC 20/224, triangle) are displayed. Values for 1:100 dilutions of sample 1 for operators 2 and 3 were > ULOQ.

**Table 3.** ELISA dilution linearity.

| IU/mL values over 1:100 to 1:1,600 dilution series |  |  |  |  |  |  |  |  |  |  |  |
| --- | --- | --- | --- | --- | --- | --- | --- | --- | --- | --- | --- |
| Sample | Operator 1 |  |  | Operator 2 |  |  | Operator 3 |  |  | All values |  |
|  | Mean | R-squared | %CV | Mean | R-squared | %CV | Mean | R-squared | %CV | Mean | %CV |
| NIBSC 20/226 | 2020.8 | 0.9995 | 3.2 | 2128.1 | 0.9951 | 6.3 | 1664 | 0.9926 | 20.5 | 1916.6 | 15.6 |
| NIBSC-20/228 | 942 | 0.8433 | 20.2 | 983.4 | 0.9996 | 3.6 | 743.8 | 0.9887 | 13.4 | 889.7 | 17.9 |
| NIBSC-20/244 | 1024.9 | 0.9905 | 6.7 | 1030.2 | 0.9905 | 6.2 | 929.4 | 0.9934 | 12.7 | 994.8 | 9.4 |

Mean, R-squared and precision (%CV) values of anti-LASV-NP IgG IU/mL determinations across serial dilutions of anti-LASV-NP IgG positive samples tested by three operators.

#### Concordance between commercial kit and developed ELISA

Serum samples with a range of anti-LASV-NP IgG levels determined using the developed ELISA (ten negative and eleven positive samples) were retested with both the developed ELISA and commercial ReLASV® Pan-Lassa NP IgG/IgM ELISA Kit, to evaluate concordance in assay specificity between these two methods. Each sample was tested by two operators in both assays. 100% concordance was observed for positive and negative determinations between the two assays across both operators (Table 4). A direct comparison of quantitative outputs of the two assays was not possible as the commercial kit reports anti-LASV-NP IgG values in μg/mL and the developed assay in IU/mL. However, there was a strong and significant positive correlation between these positive sample values for the two assays (r = 0.964, p <0.0001, non-parametric Spearman correlation) (S1 Fig).

**Table 4.** Concordance between commercial and developed ELISAs.

| Serum Sample | Commercial ELISA kit |  |  |  |  | Developed in-house ELISA |  |  |  |  |
| --- | --- | --- | --- | --- | --- | --- | --- | --- | --- | --- |
|  | Operator 1 |  | Operator 2 |  | Mean IgG | Operator 1 |  | Operator 2 |  | Mean IgG |
|  | Mean OD | Result | Mean OD | Result | µg/mL | Mean OD | Result | Mean OD | Result | IU/mL |
| NS-01 | 0.008 | Neg<sup>a</sup> | 0.007 | Neg<sup>a</sup> | NA<sup>c</sup> | 0.028 | Neg<sup>a</sup> | 0.070 | Neg<sup>a</sup> | NA<sup>c</sup> |
| NS-02 | 0.012 | Neg<sup>a</sup> | 0.011 | Neg<sup>a</sup> | NA<sup>c</sup> | 0.024 | Neg<sup>a</sup> | 0.039 | Neg<sup>a</sup> | NA<sup>c</sup> |
| NS-03 | 0.017 | Neg<sup>a</sup> | 0.018 | Neg<sup>a</sup> | NA<sup>c</sup> | 0.053 | Neg<sup>a</sup> | 0.075 | Neg<sup>a</sup> | NA<sup>c</sup> |
| NS-04 | 0.017 | Neg<sup>a</sup> | 0.016 | Neg<sup>a</sup> | NA<sup>c</sup> | 0.050 | Neg<sup>a</sup> | 0.058 | Neg<sup>a</sup> | NA<sup>c</sup> |
| NS-05 | 0.020 | Neg<sup>a</sup> | 0.021 | Neg<sup>a</sup> | NA<sup>c</sup> | 0.018 | Neg<sup>a</sup> | 0.025 | Neg<sup>a</sup> | NA<sup>c</sup> |
| NS-06 | 0.009 | Neg<sup>a</sup> | 0.012 | Neg<sup>a</sup> | NA<sup>c</sup> | 0.023 | Neg<sup>a</sup> | 0.033 | Neg<sup>a</sup> | NA<sup>c</sup> |
| NS-07 | 0.008 | Neg<sup>a</sup> | 0.007 | Neg<sup>a</sup> | NA<sup>c</sup> | 0.013 | Neg<sup>a</sup> | 0.024 | Neg<sup>a</sup> | NA<sup>c</sup> |
| NS-08 | 0.019 | Neg<sup>a</sup> | 0.019 | Neg<sup>a</sup> | NA<sup>c</sup> | 0.025 | Neg<sup>a</sup> | 0.029 | Neg<sup>a</sup> | NA<sup>c</sup> |
| NS-09 | 0.027 | Neg<sup>a</sup> | 0.025 | Neg<sup>a</sup> | NA<sup>c</sup> | 0.033 | Neg<sup>a</sup> | 0.050 | Neg<sup>a</sup> | NA<sup>c</sup> |
| NS-10 | 0.014 | Neg<sup>a</sup> | 0.010 | Neg<sup>a</sup> | NA<sup>c</sup> | 0.081 | Neg<sup>a</sup> | 0.108 | Neg<sup>a</sup> | NA<sup>c</sup> |
| C105-14 | 0.666 | Pos<sup>b</sup> | 0.584 | Pos<sup>b</sup> | 2.90 | 1.984 | Pos<sup>b</sup> | 2.121 | Pos<sup>b</sup> | 4.75 |
| C105-15 | 0.707 | Pos<sup>b</sup> | 0.644 | Pos<sup>b</sup> | 3.17 | 2.193 | Pos<sup>b</sup> | 2.211 | Pos<sup>b</sup> | 5.73 |
| C105-16 | 0.473 | Pos<sup>b</sup> | 0.385 | Pos<sup>b</sup> | 1.90 | 1.696 | Pos<sup>b</sup> | 1.830 | Pos<sup>b</sup> | 3.36 |
| C105-17 | 0.295 | Pos<sup>b</sup> | 0.213 | Pos<sup>b</sup> | 1.09 | 1.571 | Pos<sup>b</sup> | 1.695 | Pos<sup>b</sup> | 2.88 |
| NIBSC-20/204 | 1.081 | Pos<sup>b</sup> | 0.863 | Pos<sup>b</sup> | 4.91 | 2.350 | Pos<sup>b</sup> | 2.443 | Pos<sup>b</sup> | 7.41 |
| NIBSC-20/222 | 0.443 | Pos<sup>b</sup> | 0.334 | Pos<sup>b</sup> | 1.70 | 1.718 | Pos<sup>b</sup> | 1.853 | Pos<sup>b</sup> | 3.45 |
| NIBSC-20/226 | 1.656 | Pos<sup>b</sup> | 1.280 | Pos<sup>b</sup> | 8.73 | 2.947 | Pos<sup>b</sup> | 2.990 | Pos<sup>b</sup> | >ULOQ<sup>d</sup> |
| NIBSC-20/228 | 1.143 | Pos<sup>b</sup> | 0.915 | Pos<sup>b</sup> | 5.29 | 2.441 | Pos<sup>b</sup> | 2.618 | Pos<sup>b</sup> | 9.01 |
| NIBSC-20/244 | 1.416 | Pos<sup>b</sup> | 1.339 | Pos<sup>b</sup> | 7.95 | 2.496 | Pos<sup>b</sup> | 3.034 | Pos<sup>b</sup> | 8.96 |
| NIBSC-20/246 | 0.367 | Pos<sup>b</sup> | 0.294 | Pos<sup>b</sup> | 1.44 | 1.492 | Pos<sup>b</sup> | 1.528 | Pos<sup>b</sup> | 2.49 |
| NIBSC-20/248 | 0.277 | Pos<sup>b</sup> | 0.210 | Pos<sup>b</sup> | 1.04 | 1.154 | Pos<sup>b</sup> | 1.164 | Pos<sup>b</sup> | 1.60 |

Full concordance in anti-LASV-NP IgG ELISA results between the commercial and developed ELISAs. Samples were tested in both assays by two operators.
<sup>a</sup>Neg: negative result below assay positivity point. <sup>b</sup>Pos: positive result above assay positivity point. <sup>c</sup>NA: not applicable (negative samples). <sup>d</sup>ULOQ: upper limit of quantification.

#### Comparison of manual and automated ELISA plate washing

ELISA plates may be washed manually using a multichannel pipettor or by using an automated plate washer, with such a difference being a potential source of assay variation. To assess if the manner of plate washing impacted on assay outcome, three operators each tested eleven negative and ten positive anti-LASV-NP IgG serum samples, with samples duplicated across two ELISA plates. One plate was manually washed and the other using an automated plate washer (Tecan). All OD values for the eleven negative samples tested by three operators were below the assay positivity point of 0.182 for manual (mean OD = 0.040, median = 0.035) and automated plate washing (mean OD = 0.039, median = 0.030) (data not shown). Fig 7 and Table 5 display data for ten positive samples in IU/mL. Data for one sample above the assay ULOQ were excluded from further analyses. There was no significant difference in IU/mL values for manual (mean = 4.87, median = 5.01) or automated (mean = 5.52, median 5.74) plate washing (p = 0.1042; Wilcoxon matched pairs signed rank test). Data were precise across the three operators and plate washing method with nine of ten (90%) of data having CVs of <25%

**Fig 7.**
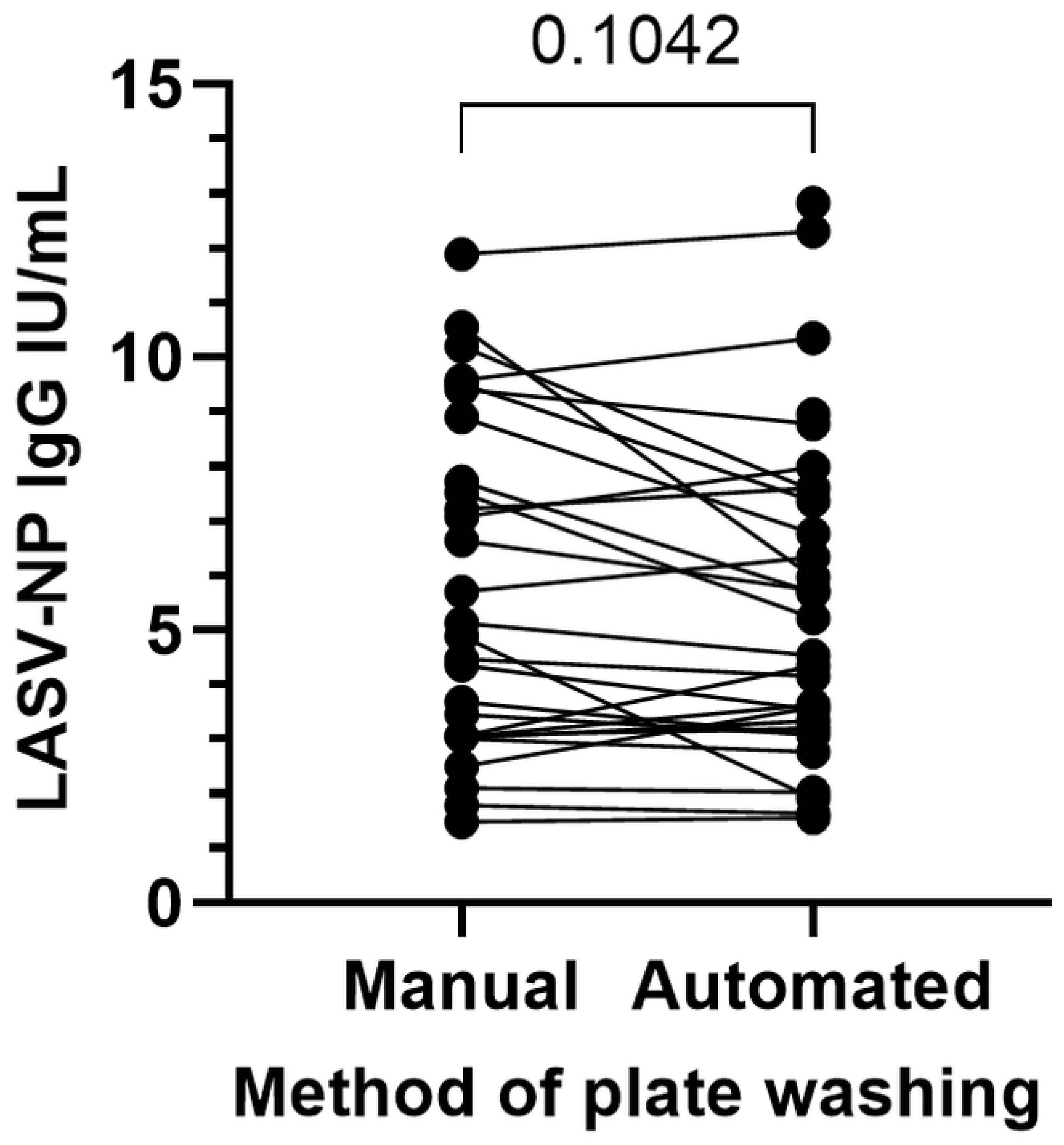
Comparison of manual and automated plate washing. Comparison of anti-LASV-NP IgG IU/mL results for positive samples tested in ELISA plates washed manually or with an automated plate washer. The p value of a Wilcoxon matched pairs signed rank test for the two datasets is displayed above the datapoints.

**Table 5.** Comparison of manual and automated ELISA plate washing.

| Serum Sample | Operator 1 IU/mL |  | Operator 2 IU/mL |  | Operator 3 IU/mL |  | Mean of operators |  | All data |  |
| --- | --- | --- | --- | --- | --- | --- | --- | --- | --- | --- |
|  | Manual | Automated | Manual | Automated | Manual | Automated | Manual | Automated | Mean | %CV |
| C105-18 | 5.70 | 6.34 | 6.64 | 5.72 | 7.51 | 5.21 | 6.62 | 5.76 | 6.19 | 13.3 |
| C105-19 | 7.72 | 5.69 | 8.90 | 6.77 | 10.55 | 5.97 | 9.05 | 6.14 | 7.60 | 24.6 |
| C105-20 | 3.04 | 3.62 | 4.47 | 4.15 | 2.49 | 3.57 | 3.33 | 3.78 | 3.56 | 20.2 |
| C105-21 | 3.02 | 3.33 | 4.35 | 3.54 | 3.06 | 3.20 | 3.48 | 3.36 | 3.42 | 14.5 |
| NIBSC-20/204 | 7.07 | 7.98 | 9.41 | 8.77 | 10.19 | 7.56 | 8.89 | 8.10 | 8.49 | 13.9 |
| NIBSC-20/222 | 3.05 | 4.33 | 5.12 | 4.53 | 3.68 | 3.07 | 3.95 | 3.98 | 3.97 | 21.2 |
| NIBSC-20/226 | 11.74 | >ULOQ<sup>a</sup> | >ULOQ<sup>a</sup> | >ULOQ<sup>a</sup> | >ULOQ<sup>a</sup> | >ULOQ<sup>a</sup> | N/A<sup>b</sup> | N/A<sup>b</sup> | N/A<sup>b</sup> | N/A<sup>b</sup> |
| NIBSC-20/228 | 7.20 | 7.60 | 9.57 | 10.35 | >ULOQ | 8.95 | 8.39 | 8.97 | 8.74 | 15.1 |
| NIBSC-20/244 | 9.50 | 7.37 | 11.88 | 12.30 | >ULOQ | 12.82 | 10.69 | 10.83 | 10.77 | 21.3 |
| NIBSC-20/246 | 4.88 | 1.91 | 3.44 | 3.08 | 3.00 | 2.77 | 3.78 | 2.59 | 3.18 | 30.8 |
| NIBSC-20/248 | 1.48 | 1.56 | 2.11 | 2.03 | 1.78 | 1.63 | 1.79 | 1.74 | 1.76 | 14.6 |

Comparison of anti-LASV-NP IgG IU/mL results for ELISA plates washed manually or with an automated plate washer.
<sup>a</sup>ULOQ: upper limit of quantification. <sup>b</sup>N/A: not applicable

#### Anti-LASV-NP IgG stability over serum freeze-thaw cycles

Five anti-LASV-NP IgG positive serum samples were aliquoted and subjected to one to five freeze-thaw cycles (frozen at -80°C for at least one hour and thawed and held at ambient temperature for least one hour). Freeze-thaw aliquots were simultaneously assessed for anti-LASV-NP IgG IU/mL content. Fig 8 demonstrates that IU/mL values were stable with no significant differences in values over the five freeze-thaw cycles (p = 0.578, non-parametric paired Friedman test).

**Fig 8.**
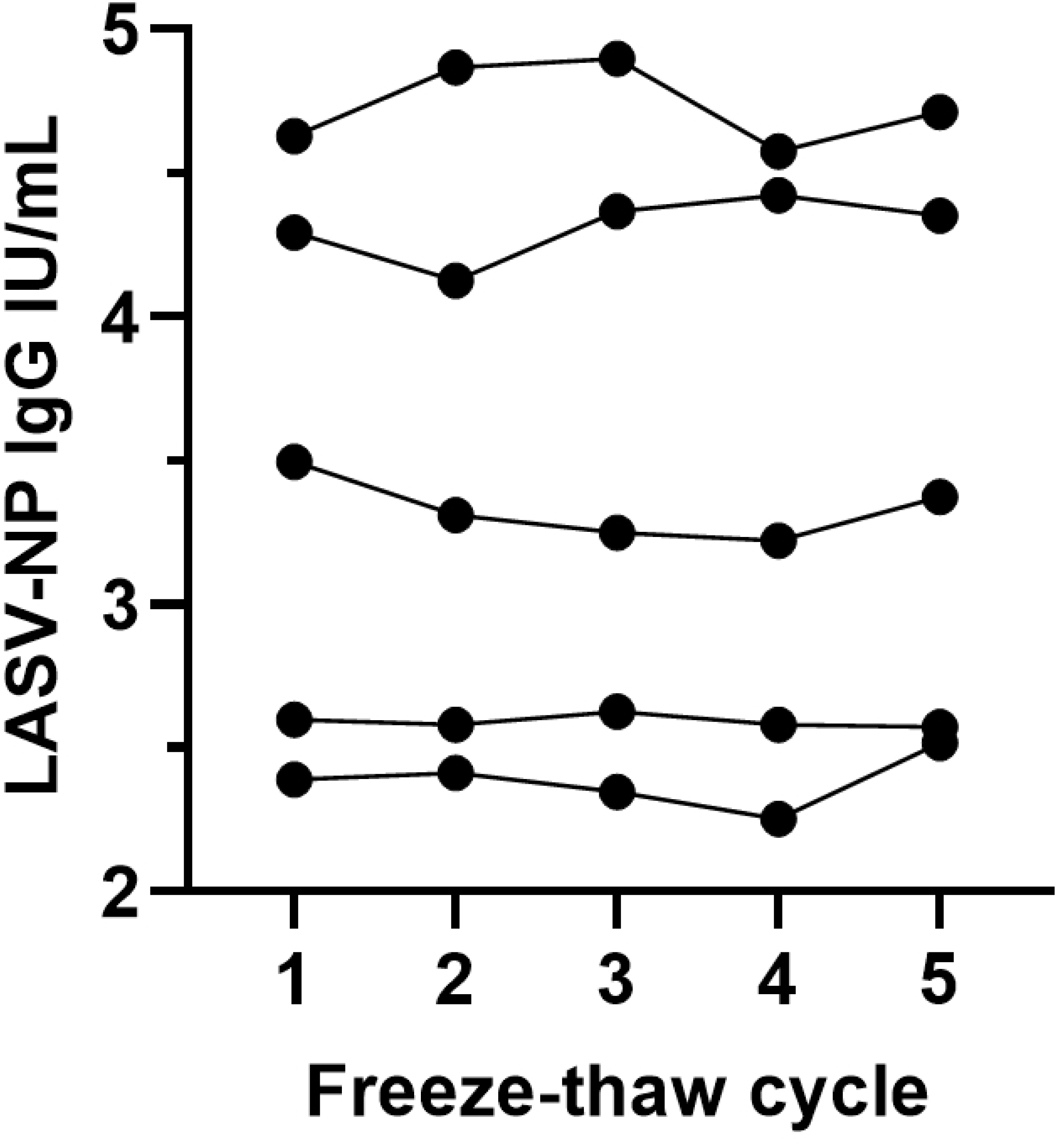
Freeze-thaw stability. Stability of anti-LASV-NP IgG IU/mL values in five anti-LASV-NP IgG positive serum samples over 1 to 5 freeze-thaw cycles between -80°C and room temperature.

#### Assay selectivity in hemolyzed serum samples

A small number of clinical trial serum samples may originate from whole blood samples that have undergone hemolysis, releasing red blood cell contents that may affect ELISA selectivity: the ability of the assay to detect the analyte in the presence of potentially inferring serum components. Hemolyzed human serum samples were obtained from individuals with no known exposure to LASV and shown to be negative for anti-LASV-NP IgG using the developed ELISA, with ODs between 0 and 0.140, below the assay positivity point of OD 0.182. Five anti-LASV-NP IgG positive samples were spiked into hemolyzed serum and tested on four occasions by two operators. This assay was considered selective if anti-LASV-NP IgG values for positive serum samples spiked into hemolyzed serum were between 80% and 120% of values for positive sera in the absence of hemolyzed serum. Assay selectivity in the presence of hemolyzed serum was demonstrated by the overall mean recovery of 87.3% (range 70.4% to 106.1%) of anti-LASV-NP IgG IU/mL relative to un-hemolyzed serum samples. Recoveries between 80% and 120% were achieved for all five positive samples with this achieved in seventeen out of twenty (85%) occasions across the replicate tests (Table 6).

**Table 6.** ELISA selectivity in hemolyzed serum.

| LASV-NP IgG positive<br>serum sample | % recovery of LASV-NP IgG spiked in to hemolyzed serum |  |  |  |  |
| --- | --- | --- | --- | --- | --- |
|  | Run 1 | Run 2 | Run 3 | Run 4 | Mean |
| C105-22 | 99.4 | 78.8 | 90.4 | 85.4 | 88.5 |
| C105-23 | 95.4 | 84.6 | 81.0 | 86.1 | 86.8 |
| C105-24 | 75.3 | 106.1 | 90.0 | 96.3 | 91.9 |
| C105-25 | 91.1 | 88.8 | 93.9 | 90.8 | 91.1 |
| C105-26 | 80.2 | 81.1 | 70.4 | 81.0 | 78.2 |
| Mean | 88.3 | 87.9 | 85.1 | 87.9 | 87.3 |

Per cent recovery of anti-LASV-NP IgG values for positive serum samples in the presence of hemolyzed serum compared to un-hemolyzed serum.

#### ELISA assay and test sample acceptance criteria and data analysis

ELISA plate and test sample acceptance criteria were established based on the collated qualification data of this study. Table 7 displays means, standard deviations and %CVs for reagent blank and negative control OD values and LPC and HPC IU/mL values along with information regarding the standard curves. The difference between the standard curve upper (D) and lower (A) asymptote interpolated IU/mL values of the 4-PL fitting curve and R-squared values as a measure of good fit to the curve are reported. For an ELISA plate result to be accepted, the reagent blank OD must be <0.1, negative control OD must be <0.182 and low and high positive controls must have recoveries of 75% to 125% of expected anti-LASV-NP IgG IU/mL values (0.349 ≤LPC ≤ 0.581 and 1.394 ≤HPC ≤2.323). The standard curve must have an R-squared value > 0.98 with the difference between the upper (D) and lower (A) asymptote interpolated IU/mL values (D-A) being > 1.895, which equates to the mean of these values minus three standard deviations. If any of these assay acceptance criteria are not met, the whole plate must be repeated.

**Table 7.**
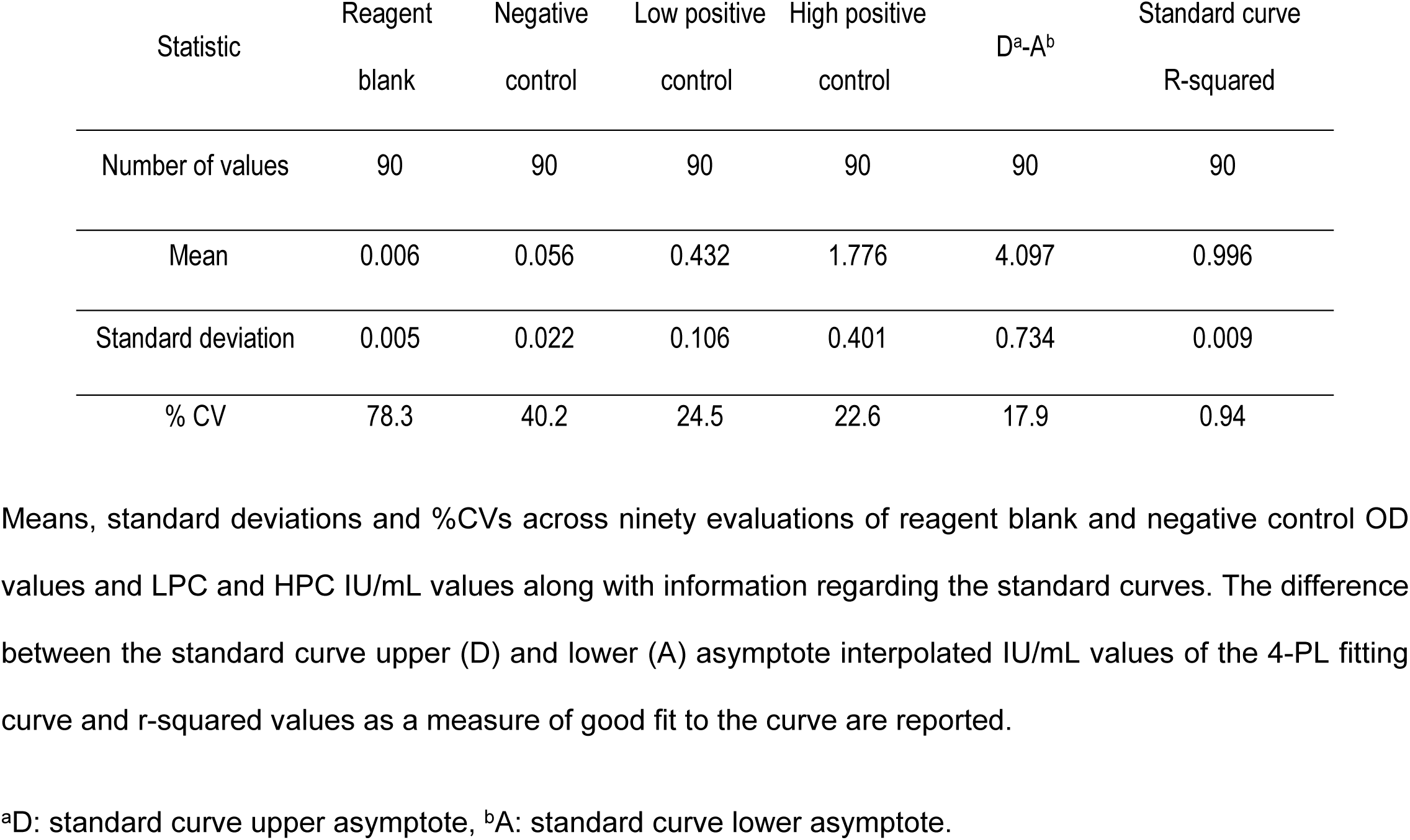
Collated ELISA qualification data for assay reagent blank, controls and standard curves.

OD values across replicate wells for individual standard curve and positive controls data points must have CVs ≤25%. All test samples within an ELISA plate that does not meet these criteria warrant repeat testing. Within an acceptable ELISA plate, one of three replicate wells of an individual positive test sample can be excluded if OD values have a CV >25%. The sample would warrant repeat testing if the CV is still >25%. All test samples within one ELISA plate warrant repeat testing if more than four individual samples show CVs >25%. Replicate wells for standard curve, negative controls and reagent blank wells cannot be excluded.

The assay dynamic range extends from 0.121 IU/mL: the LLOQ, to 7.744 IU/mL: the ULOQ. Samples producing OD values below the established positivity point of 0.182 at a 1:100 dilution are classified as negative and assigned an arbitrary concentration value of 1 IU/mL for data analysis. Samples with OD values above 0.182 are classified as positive. If a positive sample generates a signal above the positivity point but below the LLOQ at a 1:100 dilution, a final concentration of 5 IU/mL is assigned for data analysis. Samples exceeding the ULOQ (7.744 IU/mL) must be retested at a higher dilution. Final anti-LASV NP IgG concentrations should incorporate the dilution factor used for testing. Table S5 provides a summary of all assay acceptance criteria.

## Conclusions

ELISA is a vital tool for measuring antibodies against LASV-NP, enabling sensitive and specific detection of IgG responses targeting this antigen. Since NP is highly immunogenic [17–19], antibody detection provides critical information for diagnosing infection, assessing immunity and conducting sero-epidemiological studies. This assay is essential for monitoring population exposure, guiding public health interventions and supporting vaccine development against LASV.

We report the successful development and qualification of an anti-LASV-NP IgG ELISA entirely at a primary immunology laboratory in Ghana. This represents a key milestone for Lassa fever research in West Africa, as it demonstrates that complex immunogenicity endpoint assays can be established and performed in endemic regions, reducing reliance on distant reference laboratories. The need for development of such an assay arose with the discontinuation of the commercially available LASV® Pan-Lassa NP IgG/IgM ELISA Kit. The developed ELISA and commercial kit were shown to have equivalent performance in qualitative detection of anti-LASV-NP IgG, with the two assays having a strong positive correlation in quantification, although with different reported units of IgG concentration (IU/mL and μg/mL, respectively).

The developed ELISA utilizing LASV-NP lineage IV as coating antigen allowed detection of IgG reactive to NP from the different LASV lineages predominating across West Africa, presumably due to the relatively high level of shared amino acid sequence between LASV-NP lineages [5]. Screening of clinical trial samples to detect serum IgG reactivity to NP from multiple LASV lineages is facilitated in terms of cost, time and sample volumes with one ELISA utilizing lineage IV NP as coating antigen. Assay qualification demonstrated a reproducible assay with acceptable sensitivity, specificity, selectivity and precision. Clear discrimination in ELISA signal was demonstrated between serum samples from subjects previously exposed to LASV and those with no exposure history. The assay was shown to be reliable and reproducible when performed by different operators, with precise low %CV values that were accurate to expected IgG standard concentrations over the assay dynamic range of 0.12 to 7.74 IU/mL. Linear concentrations of IgG were determined over a typical serum sample dilution range. Stable anti-LAV-NP IgG concentrations were detected over repeated freeze-thaw cycles and when employing either manual or automated ELISA plate washing.

The creation of a new LASV reference serum pool calibrated to the WHO International Standard ensures sustainability for future clinical trials and sero-epidemiological studies. This ELISA provides a standardized, reproducible and multi-lineage compatible tool for assessing anti-LASV-NP IgG responses and monitoring vaccine-induced or natural immunity. By establishing this capability locally, the study enhances the ability of West African laboratories to conduct high-quality immunological assessments in populations most affected by Lassa fever and other pathogens.

## Acknowledgements

We thank the CEPI and EDCTP project teams for their continuous support and helpful discussions. We thank volunteers whose samples were used in this study and the following persons for supporting roles in this work: Bridgette Connell, Hester Kuipers, Jennifer Lehrman, Kara Bickham, Gretchen Meller, Thomas Postler, Michelle Lam, Samson Ndakala, Natalia Fernandez, Suzanna Francis, Sadou Sako (all from IAVI) and Theophilus Brenko (NMIMR, Ghana). We thank Luis Branco, Doug Simson and Zalgen Labs for providing recombinant LASV proteins.

## Funding

This work was funded by Coalition for Epidemic Preparedness Innovations (CEPI) (award reference number: 276869) and the EDCTP2 programme supported by the European Union EDCTP (grant number: RIA2019LV-3053).

## Supporting information

**S1 Table.** Standard curve OD values and positive control concentrations.

|  | LASV-NP coating concentration (total protein) and lineage |  |  |  |
| --- | --- | --- | --- | --- |
| Nonlinear Curve Fit | 1 µg/mL<br>IV | 1 µg/mL<br>II/III/IV | 2 µg/mL<br>IV | 2 µg/mL<br>II/III/IV |
| Bottom | 0.050 | 0.040 | 0.180 | 0.999 |
| Top | 2.417 | 1.843 | 2.830 | 2.744 |
| IC50 | 1.732 | 1.727 | 2.760 | 7.273 |
| Hill Slope | 1.096 | 1.075 | 1.006 | 1.762 |
| LogIC50 | 0.239 | 0.237 | 0.441 | 0.862 |
| Span | 2.367 | 1.803 | 2.650 | 1.744 |
| R squared | 0.932 | 0.814 | 0.996 | 0.763 |
| Interpolated IU/mL concentration of<br>Positive Control (Nominal<br>concentration: 10 IU/mL) | 4.303 | 6.207 | 8.901 | 9.220 |
| Negative Control (OD) | 0.201 | 0.058 | 0.099 | 0.098 |

**S2 Table.** New reference serum pool anti-LASV-NP IgG concentrations.

| Serum pool and<br>standard curve dilutions |  | Lineage II/III/IV<br>Mean = 417.992 IU/mL |  |  |  | Lineage IV<br>Mean = 544.847 IU/mL |  |  |  |
| --- | --- | --- | --- | --- | --- | --- | --- | --- | --- |
|  |  | Mean | 434.425 | Mean | 401.560 | Mean | 550.496 | Mean | 539.197 |
|  |  | SD | 55.507 | SD | 22.205 | SD | 29.064 | SD | 22.920 |
|  |  | CV% | 12.777 | CV% | 5.530 | CV% | 5.280 | CV% | 4.251 |
| Serum pool and<br>standard curve dilutions |  | Run 1 |  | Run 2 |  | Run 1 |  | Run 2 |  |
| Dilution<br>Factor | Pool / STD | Interpolated<br>Concentration | Result | Interpolated<br>Concentration | Result | Interpolated<br>Concentration | Result | Interpolated<br>Concentration | Result |
| 50 | Pool STD1 | 10.02 | 500.999 | 7.382 | 369.116 | 12.138 | 606.876 | 10.441 | 522.042 |
| 100 | Pool STD2 | 4.783 | 478.306 | 4.181 | 418.067 | 5.479 | 547.882 | 5.169 | 516.897 |
| 200 | Pool STD3 | 2.359 | 471.718 | 2.029 | 405.841 | 2.671 | 534.219 | 2.803 | 560.665 |
| 400 | Pool STD4 | 1.001 | 400.263 | 1.033 | 413.215 | 1.323 | 529.145 | 1.393 | 557.184 |
| 800 | Pool STD5 | 0.478 | 382.042 | 0.497 | 397.828 | 0.666 | 532.757 | 0.706 | 565.124 |
| 1600 | Pool STD6 | 0.233 | 373.221 | 0.251 | 401.795 | 0.345 | 552.099 | 0.347 | 555.942 |
| 3200 | Pool STD7 | 0.120 | 385.590 | 0.108 | 345.757 | 0.175 | 561.060 | 0.156 | 497.680 |
| 6400 | Pool STD8 | 0.066 | 420.072 | 0.045 | 288.777 | 0.089 | 567.761 | 0.064 | 409.402 |
| 12800 | Pool STD9 | 0.040 | 513.184 | 0.015 | 190.232 | 0.047 | 600.475 | 0.009 | 118.547 |
| 25600 | Pool STD10 | 0.029 | 734.687 | NA<sup>a</sup> | NA<sup>a</sup> | 0.027 | 701.632 | NA<sup>a</sup> | NA<sup>a</sup> |

Determination of interim anti-LASV-NP IgG concentrations (IU/mL) in a new reference serum pool by serial dilution of pool and WHO reference standard. The new reference serum pool was assigned values of 418.0 and 544.8 IU/mL for LASV-NP lineages II, III and IV and lineage IV alone, respectively.
<sup>a</sup>NA: not available.

**S3 Table.** Assay specificity with negative samples.

| Serum<br>sample | Operator 1 OD values |  |  | Operator 2 OD values |  |  | Operator 3 OD values |  |  | All operators |  |  |
| --- | --- | --- | --- | --- | --- | --- | --- | --- | --- | --- | --- | --- |
|  | RUN 1 | RUN 2 | RUN 3 | RUN 1 | RUN 2 | RUN 3 | RUN 1 | RUN 2 | RUN 3 | Mean | %CV | % results |
|  |  |  |  |  |  |  |  |  |  |  |  | negative<br>OD ≤0.182 |
| NS-11 | 0.124 | 0.082 | 0.116 | 0.118 | 0.075 | 0.071 | 0.106 | 0.069 | 0.062 | 0.091 | 26.7 | 100 |
| NS-12 | 0.122 | 0.105 | 0.141 | 0.077 | 0.070 | 0.067 | 0.093 | 0.082 | 0.061 | 0.091 | 29.7 | 100 |
| NS-13 | 0.097 | 0.075 | 0.075 | 0.054 | 0.046 | 0.046 | 0.059 | 0.045 | 0.033 | 0.059 | 33.8 | 100 |
| NS-14 | 0.169 | 0.140 | 0.123 | 0.100 | 0.055 | 0.058 | 0.119 | 0.057 | 0.046 | 0.096 | 46.0 | 100 |
| NS-15 | 0.131 | 0.207 | 0.223 | 0.133 | 0.115 | 0.122 | 0.125 | 0.115 | 0.129 | 0.145 | 28.1 | 77.8 |
| NS-16 | 0.091 | 0.075 | 0.061 | 0.095 | 0.045 | 0.047 | 0.071 | 0.045 | 0.031 | 0.062 | 35.7 | 100 |
| NS-17 | 0.115 | 0.077 | 0.077 | 0.080 | 0.043 | 0.048 | 0.062 | 0.044 | 0.033 | 0.064 | 39.8 | 100 |
| NS-18 | 0.117 | 0.022 | 0.071 | 0.087 | 0.049 | 0.057 | 0.072 | 0.051 | 0.042 | 0.063 | 44.0 | 100 |
| NS-19 | 0.000 | 0.157 | 0.145 | 0.119 | 0.081 | 0.070 | 0.125 | 0.091 | 0.075 | 0.096 | 49.6 | 100 |
| NS-20 | 0.146 | 0.100 | 0.132 | 0.094 | 0.082 | 0.076 | 0.128 | 0.071 | 0.066 | 0.100 | 29.4 | 100 |
| NS-21 | 0.145 | 0.186 | 0.163 | 0.155 | 0.114 | 0.123 | 0.144 | 0.110 | 0.100 | 0.138 | 20.4 | 88.9 |
| NS-22 | 0.203 | 0.132 | 0.188 | 0.144 | 0.101 | 0.132 | 0.144 | 0.092 | 0.081 | 0.135 | 30.5 | 77.8 |
| NS-23 | 0.240 | 0.050 | 0.057 | 0.049 | 0.026 | 0.051 | 0.054 | 0.030 | 0.039 | 0.066 | 99.5 | 100 |
| NS-24 | 0.026 | 0.025 | 0.018 | 0.018 | 0.010 | 0.012 | 0.023 | 0.012 | 0.003 | 0.016 | 48.0 | 100 |
| NS-25 | 0.136 | 0.087 | 0.098 | 0.067 | 0.047 | 0.049 | 0.103 | 0.066 | 0.060 | 0.079 | 36.9 | 100 |
| NS-26 | 0.115 | 0.135 | 0.102 | 0.071 | 0.062 | 0.061 | 0.088 | 0.072 | 0.052 | 0.084 | 33.2 | 100 |
| NS-27 | 0.030 | 0.029 | 0.025 | 0.013 | 0.012 | 0.012 | 0.020 | 0.017 | 0.003 | 0.018 | 51.0 | 100 |
| NS-28 | 0.039 | 0.031 | 0.027 | 0.024 | 0.020 | 0.020 | 0.032 | 0.018 | 0.009 | 0.025 | 35.6 | 100 |
| NS-29 | 0.166 | 0.140 | 0.124 | 0.116 | 0.083 | 0.081 | 0.132 | 0.078 | 0.068 | 0.110 | 30.6 | 100 |
| NS-30 | 0.061 | 0.035 | 0.049 | 0.035 | 0.025 | 0.041 | 0.037 | 0.025 | 0.034 | 0.038 | 30.3 | 100 |
| NS-31 | 0.070 | 0.059 | 0.053 | 0.052 | 0.034 | 0.049 | 0.053 | 0.033 | 0.027 | 0.048 | 29.2 | 100 |
| NS-32 | 0.074 | 0.076 | 0.069 | 0.057 | 0.031 | 0.048 | 0.051 | 0.063 | 0.050 | 0.058 | 25.2 | 100 |
| NS-33 | 0.041 | 0.042 | 0.067 | 0.042 | 0.019 | 0.032 | 0.038 | 0.038 | 0.037 | 0.040 | 31.6 | 100 |
| NS-34 | 0.151 | 0.152 | 0.187 | 0.189 | 0.080 | 0.148 | 0.113 | 0.087 | 0.101 | 0.134 | 30.5 | 77.8 |
| NS-35 | 0.102 | 0.094 | 0.105 | 0.097 | 0.045 | 0.074 | 0.067 | 0.063 | 0.071 | 0.080 | 25.9 | 100 |
| NS-36 | 0.051 | 0.087 | 0.098 | 0.040 | 0.019 | 0.046 | 0.037 | 0.036 | 0.060 | 0.053 | 47.8 | 100 |
| NS-37 | 0.044 | 0.048 | 0.056 | 0.047 | 0.020 | 0.031 | 0.025 | 0.032 | 0.031 | 0.037 | 32.3 | 100 |
| NS-38 | 0.115 | 0.100 | 0.143 | 0.073 | 0.037 | 0.065 | 0.051 | 0.063 | 0.066 | 0.079 | 42.5 | 100 |
| NS-39 | 0.073 | 0.087 | 0.086 | 0.067 | 0.045 | 0.066 | 0.057 | 0.066 | 0.056 | 0.067 | 20.4 | 100 |
| NS-40 | 0.044 | 0.055 | 0.046 | 0.037 | 0.022 | 0.027 | 0.025 | 0.023 | 0.021 | 0.033 | 37.5 | 100 |
| NS-41 | 0.052 | 0.049 | 0.057 | 0.034 | 0.022 | 0.024 | 0.026 | 0.030 | 0.031 | 0.036 | 36.0 | 100 |
| NS-42 | 0.039 | 0.029 | 0.059 | 0.026 | 0.014 | 0.019 | 0.018 | 0.019 | 0.018 | 0.027 | 53.3 | 100 |
| NS-43 | 0.025 | 0.043 | 0.050 | 0.030 | 0.020 | 0.024 | 0.026 | 0.032 | 0.043 | 0.032 | 32.2 | 100 |
| NS-44 | 0.043 | 0.072 | 0.080 | 0.060 | 0.028 | 0.032 | 0.040 | 0.103 | 0.048 | 0.056 | 44.0 | 100 |
| NS-45 | 0.189 | 0.138 | 0.212 | 0.106 | 0.047 | 0.098 | 0.112 | 0.068 | 0.114 | 0.121 | 43.8 | 77.8 |
| NS-46 | 0.065 | 0.044 | 0.055 | 0.043 | 0.021 | 0.038 | 0.038 | 0.039 | 0.049 | 0.044 | 28.4 | 100 |
| NS-47 | 0.228 | 0.228 | 0.237 | 0.184 | 0.069 | 0.157 | 0.124 | 0.094 | 0.137 | 0.162 | 37.9 | 55.6 |
| NS-48 | 0.180 | 0.172 | 0.186 | 0.146 | 0.074 | 0.149 | 0.112 | 0.127 | 0.138 | 0.143 | 24.9 | 88.9 |
| NS-49 | 0.087 | 0.078 | 0.099 | 0.065 | 0.012 | 0.048 | 0.042 | 0.039 | 0.060 | 0.059 | 45.6 | 100 |
| NS-50 | 0.055 | 0.052 | 0.060 | 0.043 | 0.020 | 0.040 | 0.029 | 0.029 | 0.039 | 0.041 | 32.2 | 100 |
| NS-51 | 0.094 | 0.145 | 0.138 | 0.073 | 0.028 | 0.080 | 0.055 | 0.056 | 0.076 | 0.083 | 46.2 | 100 |
| NS-52 | 0.175 | 0.096 | 0.155 | 0.080 | 0.063 | 0.054 | 0.043 | 0.049 | 0.078 | 0.088 | 53.1 | 100 |
| NS-53 | 0.084 | 0.084 | 0.072 | 0.055 | 0.045 | 0.036 | 0.035 | 0.035 | 0.054 | 0.056 | 36.1 | 100 |
| NS-54 | 0.023 | 0.022 | 0.024 | 0.014 | 0.009 | 0.011 | 0.025 | 0.010 | 0.013 | 0.017 | 38.9 | 100 |
| NS-55 | 0.062 | 0.060 | 0.064 | 0.036 | 0.030 | 0.032 | 0.034 | 0.025 | 0.040 | 0.043 | 35.7 | 100 |
| NS-56 | 0.079 | 0.088 | 0.082 | 0.052 | 0.059 | 0.038 | 0.043 | 0.041 | 0.044 | 0.058 | 33.4 | 100 |
| NS-57 | 0.062 | 0.056 | 0.099 | 0.036 | 0.025 | 0.052 | 0.027 | 0.026 | 0.037 | 0.047 | 51.3 | 100 |
| NS-58 | 0.263 | 0.206 | 0.285 | 0.136 | 0.126 | 0.095 | 0.020 | 0.087 | 0.145 | 0.151 | 56.6 | 66.7 |
| NS-59 | 0.127 | 0.101 | 0.110 | 0.062 | 0.067 | 0.056 | 0.023 | 0.046 | 0.060 | 0.072 | 46.2 | 100 |
| NS-60 | 0.132 | 0.117 | 0.139 | 0.068 | 0.059 | 0.055 | 0.036 | 0.046 | 0.071 | 0.080 | 48.1 | 100 |
| NS-61 | 0.055 | 0.052 | 0.071 | 0.032 | 0.022 | 0.026 | 0.047 | 0.021 | 0.026 | 0.039 | 45.7 | 100 |
| NS-62 | 0.055 | 0.052 | 0.047 | 0.030 | 0.022 | 0.026 | 0.057 | 0.023 | 0.027 | 0.038 | 39.2 | 100 |
| NS-63 | 0.043 | 0.042 | 0.047 | 0.020 | 0.020 | 0.014 | 0.050 | 0.014 | 0.018 | 0.030 | 51.6 | 100 |
| NS-64 | 0.036 | 0.022 | 0.038 | 0.015 | 0.013 | 0.020 | 0.007 | 0.014 | 0.017 | 0.020 | 52.1 | 100 |
| NS-65 | 0.138 | 0.134 | 0.202 | 0.103 | 0.078 | 0.088 | 0.029 | 0.055 | 0.097 | 0.103 | 49.5 | 88.9 |
| NS-66 | 0.127 | 0.123 | 0.136 | 0.078 | 0.076 | 0.063 | 0.034 | 0.053 | 0.075 | 0.085 | 41.8 | 100 |
| NS-67 | 0.122 | 0.100 | 0.153 | 0.082 | 0.072 | 0.073 | 0.006 | 0.050 | 0.063 | 0.080 | 52.6 | 100 |
| NS-68 | 0.093 | 0.086 | 0.073 | 0.045 | 0.035 | 0.043 | 0.010 | 0.041 | 0.048 | 0.053 | 50.0 | 100 |
| NS-69 | 0.124 | 0.095 | 0.114 | 0.067 | 0.058 | 0.063 | 0.037 | 0.071 | 0.080 | 0.079 | 35.3 | 100 |
| NS-70 | 0.133 | 0.108 | 0.089 | 0.054 | 0.048 | 0.046 | 0.019 | 0.035 | 0.069 | 0.067 | 55.0 | 100 |
| NS-71 | 0.206 | 0.199 | 0.197 | 0.134 | 0.115 | 0.098 | 0.018 | 0.073 | 0.104 | 0.127 | 50.3 | 66.7 |

Assay specificity demonstrated by anti-LASV-NP IgG ELISA OD values for sixty-one negative samples across nine tests performed by three operators. Negative values are defined as having an OD of $\leq 0.182$ . For the assay to be deemed to be specific, at least eight (88.9%) of the nine determinations for each sample must be concordant with this achieved in at least 80% of samples. In this case, determinations for fifty-four of sixty-one (88.5%) samples were concordant.

**S4 Table.** Assay specificity with positive samples.

| Serum sample | Operator 1 OD values |  |  | Operator 2 OD values |  |  | Operator 3 OD values |  |  | All operators |  |  |
| --- | --- | --- | --- | --- | --- | --- | --- | --- | --- | --- | --- | --- |
|  | RUN 1 | RUN 2 | RUN 3 | RUN 1 | RUN 2 | RUN 3 | RUN 1 | RUN 2 | RUN 3 | Mean<br>OD | % CV | % results |
|  |  |  |  |  |  |  |  |  |  |  |  | positive<br>OD >0.182 |
| NIBSC-20/226 | 3.47 | 3.40 | 2.57 | 3.52 | 3.30 | 3.12 | 3.55 | 2.78 | 3.48 | 3.24 | 10.82 | 100 |
| NIBSC-20/244 | 2.98 | 3.34 | 2.65 | 3.05 | 3.14 | 2.91 | 3.30 | 2.32 | 3.40 | 3.01 | 11.68 | 100 |
| NIBSC-20/228 | 3.04 | 3.00 | 2.42 | 3.08 | 3.15 | 2.76 | 3.21 | 2.38 | 3.11 | 2.91 | 10.76 | 100 |
| NIBSC-20/204 | 3.04 | 2.88 | 2.30 | 3.19 | 3.06 | 2.57 | 3.09 | 2.12 | 3.04 | 2.81 | 13.74 | 100 |
| C105-27 | 2.90 | 2.66 | 2.03 | 2.93 | 2.78 | 2.52 | 2.96 | 1.97 | 2.98 | 2.64 | 14.81 | 100 |
| C105-28 | 2.79 | 2.84 | 2.16 | 2.72 | 2.98 | 2.45 | 3.04 | 1.74 | 2.98 | 2.63 | 16.64 | 100 |
| C105-29 | 2.28 | 2.52 | 1.71 | 2.26 | 2.63 | 1.86 | 2.57 | 1.37 | 2.45 | 2.18 | 20.09 | 100 |
| NIBSC-20/222 | 2.47 | 2.28 | 1.68 | 2.37 | 2.32 | 1.75 | 2.44 | 1.65 | 2.24 | 2.13 | 15.94 | 100 |
| C105-30 | 2.09 | 1.98 | 1.60 | 2.28 | 2.36 | 1.83 | 2.31 | 1.24 | 2.04 | 1.97 | 18.54 | 100 |
| C105-31 | 2.10 | 2.27 | 1.39 | 1.88 | 2.18 | 1.59 | 2.41 | 1.54 | 2.19 | 1.95 | 18.66 | 100 |
| NIBSC-20/246 | 1.90 | 1.98 | 1.39 | 1.89 | 2.14 | 1.60 | 2.49 | 1.34 | 1.98 | 1.86 | 19.63 | 100 |
| C105-32 | 1.95 | 1.81 | 1.21 | 1.50 | 1.96 | 1.46 | 2.56 | 0.98 | 1.93 | 1.71 | 27.74 | 100 |
| C105-33 | 1.78 | 1.88 | 1.24 | 1.63 | 1.95 | 1.63 | 2.15 | 1.06 | 1.97 | 1.70 | 20.92 | 100 |
| C105-34 | 1.73 | 1.91 | 1.33 | 1.54 | 1.78 | 1.43 | 1.82 | 1.02 | 1.70 | 1.58 | 17.94 | 100 |
| C105-35 | 1.81 | 1.92 | 1.05 | 1.35 | 1.94 | 1.34 | 1.95 | 0.87 | 1.77 | 1.56 | 26.52 | 100 |
| C105-36 | 1.78 | 1.73 | 0.93 | 1.50 | 1.80 | 1.25 | 1.77 | 1.09 | 1.83 | 1.52 | 22.83 | 100 |
| NIBSC-20/248 | 1.58 | 1.42 | 1.09 | 1.41 | 1.79 | 1.33 | 1.74 | 1.05 | ND | 1.43 | 18.99 | 100 |
| C105-37 | 1.01 | 0.73 | 0.58 | 0.95 | 1.11 | 0.70 | 1.24 | 0.62 | 0.91 | 0.87 | 26.29 | 100 |
| C105-38 | 0.26 | 0.64 | 0.12 | 0.13 | 0.29 | 0.15 | 0.21 | 0.14 | 0.26 | 0.25 | 65.75 | 55.6 |
| C105-39 | 0.19 | 0.11 | 0.08 | 0.07 | 0.19 | 0.08 | 0.13 | 0.07 | 0.16 | 0.12 | 41.96 | 22.2 |

Assay specificity demonstrated by anti-LASV-NP IgG ELISA OD values for twenty positive samples across nine tests performed by three operators. Positive values are defined as having an OD of >0.182. For the assay to be deemed to be specific, at least eight (88.9%) of the nine determinations for each sample must be concordant with this achieved in at least 80% of samples. In this case, determinations in eighteen of twenty (90%) samples were concordant.

**S1 Fig.**
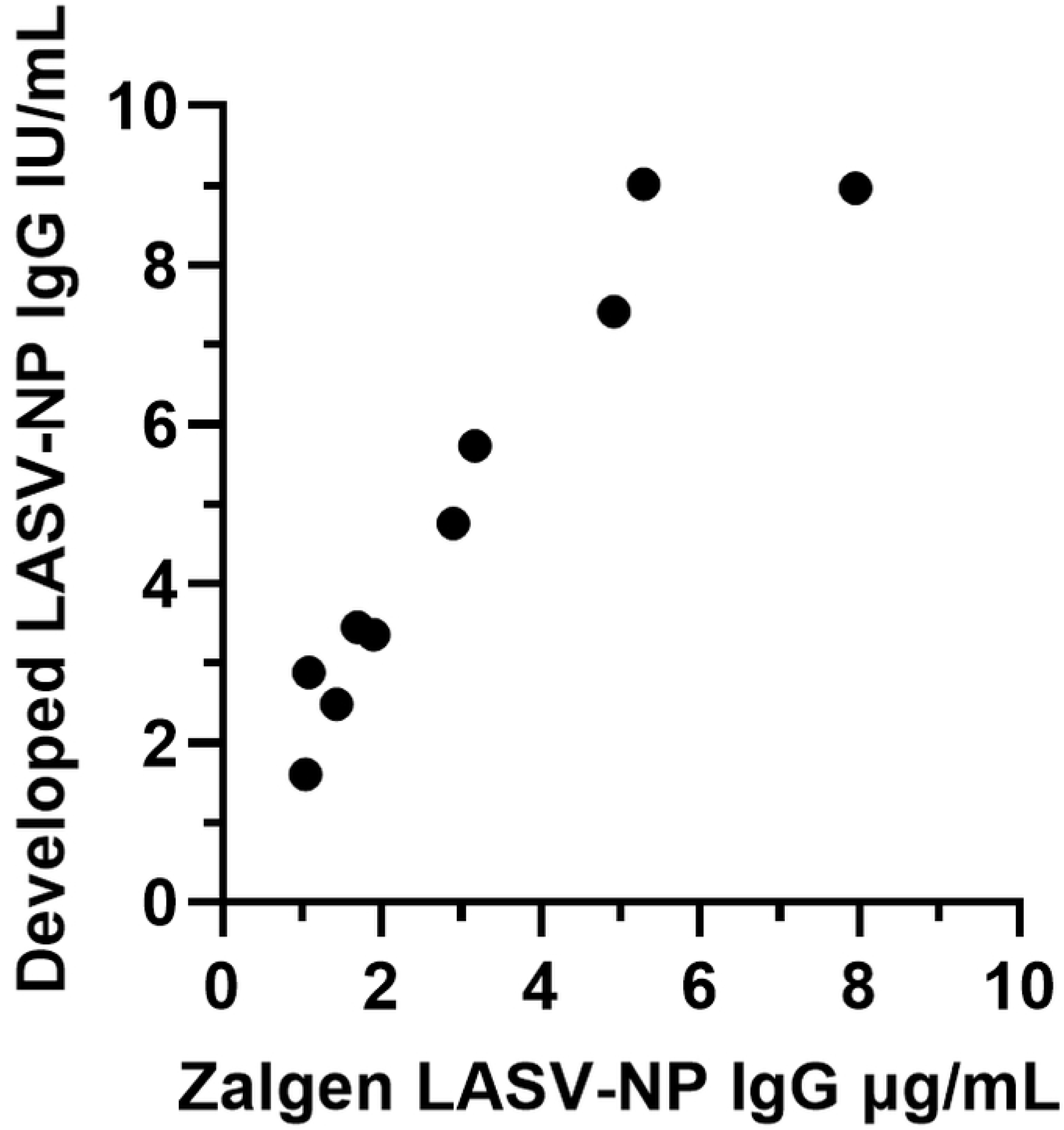
Correlation between commercial and developed ELISAs. Correlation between quantification of anti-LASV-NP IgG values determined in serum samples using a commercial Zalgen kit (expressed in μg/mL) and the developed ELISA (expressed in IU/mL). Spearman correlation demonstrates a strong and significant positive correlation between the two assays (r = 0.964, p<0.0001).

**S5 Table.** Acceptance criteria for anti-LASV-NP IgG ELISA assay

| Parameter | Acceptance criterion | Comments |
| --- | --- | --- |
| Reagent blank (OD450nm) | $\leq 0.10$ | The ELISA plate fails if this control is $>0.1$ OD |
| Assay positivity point (OD450nm) | $\leq 0.182$ | Test samples with OD of $\leq 0.182$ are deemed negative.<br>Test samples with OD $> 0.182$ are deemed positive |
| Lower limit of quantification (LLOQ)<br>(IgG IU/mL) | 0.121 | Lower limit of reliable quantification |
| Low Positive Control<br>(IgG IU/mL) | $0.349 \leq \text{LPC} \leq 0.581$ | The ELISA plate fails if values are outside of this range (75% to 125% recovery) |
| High Positive Control<br>(IgG IU/mL) | $1.394 \leq \text{HPC} \leq 2.323$ | The ELISA plate fails if values are outside of this range (75% to 125% recovery) |
| Upper limit of quantification (ULOQ)<br>(IgG IU/mL) | 7.744 | Upper limit of reliable quantification |
| CV (OD450nm) | $\leq 25 \%$ | Replicate wells of standard curve, controls and test samples must show CV values of $\leq 25\%$.<br>Does not apply to negative test samples.<br>If test sample CV $>25\%$, a total of four data points may be masked comprising:<br>• A single data point per triplicate for a maximum of four samples<br>• No data points may be masked for the standard curve, the negative controls, or the blank wells |
| Standard curve R-squared | $> 0.980$ | The ELISA plate fails if r-squared $> 0.98$ |
| 4PL standard curve<br>asymptote D - asymptote A<br>(IgG IU/mL) | $\geq 1.895$ | The ELISA plate fails if D-A $>1.895$ |

## Notes

### Competing Interest Statement

The authors have declared no competing interest.

## References

1. Garry R. Lassa fever - the road ahead. Nat Rev Microbiol. 2023;21(2):87–96. doi: 10.1038/s41579-022-00789-8.

2. Whitmer S, Strecker T, Cadar D, Dienes H, Faber K, Patel K, et al. New Lineage of Lassa Virus, Togo, 2016. Emerg Infect Dis. 2018;24(3):599–602. doi: 10.3201/eid2403.171905.

3. Yadouleton A, Picard C, Rieger T, Loko F, Cadar D, Kouthon E, et al. Lassa fever in Benin: description of the 2014 and 2016 epidemics and genetic characterization of a new Lassa virus. Emerg Microbes Infect 2020;9(1):1761–1770. doi: 10.1080/22221751.2020.1796528.

4. Aloke C, Obasi N, Aja P, Emelike C, Egwu C, Jeje O, et al. Combating Lassa Fever in West African Sub-Region: Progress, Challenges, and Future Perspectives. Viruses. 2023;15(1). doi: 10.3390/v15010146.

5. Hallam H, Hallam S, Rodriguez S, Barrett A, Beasley D, Chua A, et al. Baseline mapping of Lassa fever virology, epidemiology and vaccine research and development. NPJ Vaccines 2018;3:11. 10.1038/s41541-018-0049-5

6. WHO. Prioritizing diseases for research and development in emergency contexts. (https://www.who.int/activities/prioritizing-diseases-for-research-and-development-in-emergency-contexts#).

7. Garbutt M, Liebscher R, Wahl-Jensen V, Jones S, Moller P, Wagner R, et al. Properties of replication-competent vesicular stomatitis virus vectors expressing glycoproteins of filoviruses and arenaviruses. J Virol. 2004;78(10):5458–65. doi: 10.1128/jvi.78.10.5458-5465.2004.

8. Anderson E, Coller B. Translational success of fundamental virology: a VSV-vectored Ebola vaccine. J Virol. 2024;98(3):e0162723. doi: 10.1128/jvi.01627-23.

9. CDC. Ebola Vaccine Product Information. Centers for Disease Control and Prevention. May 1 2024 (https://www.cdc.gov/ebola/hcp/vaccines/index.html#).

10. Kallay R, Doshi RH, Muhoza P, Choi M, Legand A, Aberle-Grasse E, et al. Use of Ebola Vaccines - Worldwide, 2021-2023. MMWR Morb Mortal Wkly Rep. 2024;Apr 25;73(16):360–364. doi: 10.15585/mmwr.mm7316a1.

11. Wolf J, Jannat R, Dubey S, Troth S, Onorato M, Coller B, et al. Development of Pandemic Vaccines: ERVEBO Case Study. Vaccines (Basel). 2021;9(3). doi: 10.3390/vaccines9030190.

12. Woolsey C, Geisbert T. Current state of Ebola virus vaccines: A snapshot. PLoS Pathog. 2021;17(12):e1010078. doi: 10.1371/journal.ppat.1010078.

13. Malkin E, Zaric M, Kieh M, Baden LR, Fitz-Patrick D, Marini A, et al. Safety and Immunogenicity of an rVSV Lassa Fever Vaccine Candidate. N Engl J Med. 2025;393:1807–18. doi: 10.1056/NEJMoa2501073

14. Yun N, Walker D. Pathogenesis of Lassa fever. Viruses. 2012;Oct 9;4(10):2031–48. doi: 10.3390/v4102031. PMID: 23202452; PMCID: PMC3497040.

15. Andersen K, Shapiro B, Matranga C, Sealfon R, Lin A, Moses L, et al. Clinical Sequencing Uncovers Origins and Evolution of Lassa Virus. Cell 2015;162:738–750 doi: 10.1016/j.cell.2015.07.020

16. Bowen M, Rollin P, Ksiazek T, Hustad H, Bausch D, Demby A, et al. Genetic diversity among Lassa virus strains. Journal of Virology. 2000;74(15):6992–7004

17. Saijo M, Georges–Courbot M, Marianneau P, Romanowski V, Fukushi S, Mizutani T, et al. Development of recombinant nucleoprotein–based diagnostic systems for Lassa fever. Clinical Vaccine Immunol. 2007;14(9), 1182–1189. doi: 10.1128/CVI.00101-07.

18. Branco, L. M., Grove, J. N., Geske, F. J., Boisen, M. L., Muncy, I. J., Magliato, S. A. et al. Lassa virus–like particles displaying all major immunological determinants as a vaccine candidate for Lassa hemorrhagic fever. Virology Journal. 2010;7, 279. doi: 10.1186/1743-422X-7-279.

19. Medugu N, Adegboro B, Babazhits, M, Kadiri M, Abanida E. A review of the recent advances on Lassa fever with special reference to molecular epidemiology and progress in vaccine development. African Journal of Clinical and Experimental Microbiology. 2023;24(2), 130–146.

